# Local SNP-explained methylation variation reveals genetically anchored and exposure-associated methylation architecture in the human brain

**DOI:** 10.64898/2026.06.05.730443

**Authors:** Alexis Bennett, Elisa Kain Johnson, Nia N. Terry, Jalil Hemphill, Kynon J. M. Benjamin

**Affiliations:** Department of Psychiatry and Behavioral Sciences, Feinberg School of Medicine, Northwestern University; Weinberg College of Arts and Sciences, Northwestern University; Spelman College; McCormick School of Engineering and Applied Science, Northwestern University; Stephen M. Stahl Center for Psychiatric Neuroscience, Feinberg School of Medicine, Northwestern University · Funded by NIH Grant R00MD016964; NSF ACCESS allocation BIO250079

## Abstract

Human brain DNA methylation is shaped by inherited genetic variation and cumulative environmental experience, yet how these influences partition the methylome remains poorly resolved in postmortem cohorts with modest sample sizes and limited ancestral diversity. To map this architecture in an underrepresented population, we analyzed whole-genome bisulfite sequencing and genotype array data from 168 admixed Black American adults from the BrainSEQ consortium across three brain regions. We adapted and benchmarked SNP-based elastic-net modeling to classify variably methylated regions (VMRs) by local SNP-explained methylation variation, an approach that provided stable classification at the modest sample sizes of postmortem brain cohorts, where conventional methods are underpowered. Using this framework, we partitioned 31,143 VMRs into high and low SNP-explained classes and evaluated their generalizability in a multi-ancestry cohort of Black American and non-Hispanic white American donors. High SNP-explained VMRs were concentrated in distal intergenic sequences and, at the highest heritability levels, were enriched for H3K9me3, quiescent/repressive chromatin states and LINE/L1 elements, linking genetically anchored methylation to repeat-associated repressive chromatin across the human brain. A small subset overlapping Activity-by-Contact-defined enhancers was linked to candidate immune-related genes, including MHC class II loci. By contrast, low SNP-explained VMRs were more gene-proximal and enriched for active regulatory elements. Metadata-associated VMRs showed region-, exposure-, and donor-group-dependent enrichment across SNP-explained classes, including substance use and sociodemographic variables. Together, these findings show that the most genetically anchored component of the human brain methylome is concentrated in repressive, repeat-rich chromatin compartments involved in heterochromatin maintenance and repeat silencing, distinct from the gene-proximal, exposure-associated variation less explained by nearby SNPs. By resolving this architecture in an underrepresented population, this work clarifies how inherited variation structures the brain methylome and, given the established role of these compartments in neuronal aging, informs the interpretation of epigenomic mechanisms relevant to neuropsychiatric and neurodegenerative diseases.

## Introduction

DNA methylation (DNAm) is a stable epigenomic mark that contributes to gene regulation, cellular identity, and genome defense in the human brain, reflecting inherited genetic variation, cellular composition, and lifetime environmental experience. Twin studies estimate mean DNAm heritability at approximately 19% in blood but substantially lower levels in brain tissue (~3–10%), indicating that a large fraction of brain DNAm variation is not captured by inherited genetic effects in current study designs [1,2,3]. Therefore, distinguishing methylation variation associated with local genetic architecture from variation not explained by nearby single-nucleotide polymorphisms (SNPs) is essential for interpreting brain epigenomic studies, particularly those seeking to connect DNAm with environmental exposures and disease risk.

Large-scale methylation quantitative trait locus (meQTL) studies have shown that genetic effects on DNAm are widespread across tissues and developmental stages, including in fetal and adult brains [4,5,6]. However, how variably methylated regions (VMRs) with detectable local SNP contributions are distributed across chromatin states in the adult human brain remains incompletely resolved. Most postmortem brain DNAm studies are limited by modest sample sizes, regional heterogeneity, and cellular complexity, constraining reliable estimation of SNP-based heritability 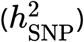 at individual methylation features. As a result, it remains unclear which VMRs show detectable local SNP contributions, how these loci are distributed across brain chromatin states, and how they differ from methylation variation less explained by nearby SNPs.

These limitations are amplified in ancestrally diverse cohorts. Most DNAm heritability studies have focused on individuals of European genetic ancestry, with African ancestry populations contributing fewer than 5% of samples to large-scale genomic studies [7,8]. This underrepresentation limits discovery and raises methodological challenges, as linkage disequilibrium (LD) structure differs across populations and can influence local SNP-based prediction. It also limits interpretation of brain methylation variation in populations that experience inequities in neurological and psychiatric disease risk, diagnosis, treatment, or outcomes, including schizophrenia [9], Alzheimer’s disease (AD) [10] and stroke [11,12]. In Black American (BA) populations, genetic ancestry, environmental exposures, and social determinants of health (SDOH) are deeply intertwined, yet brain DNAm studies have rarely had the data or analytical framework needed to separate local genetic effects from methylation variation less explained by nearby SNPs.

Here, we applied and adapted SNP-based modeling approaches to partition brain methylation variability and reveal population-relevant methylation architecture in an underrepresented cohort. Using publicly available whole-genome bisulfite sequencing (WGBS) from the BrainSEQ consortium postmortem brain tissue of 168 admixed BA adults [13,14,15], we mapped the local genetic architecture of VMRs across the hippocampus, caudate nucleus, and dorsolateral prefrontal cortex (DLPFC). Because reliable VMR-level 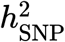 estimation is diffcult at postmortem brain sample sizes, we first established that elastic-net regression [16,17,18,19] outperforms LD-stratified genomic restricted maximum likelihood (GREML-LDMS) [20,21] in this regime, providing a stable basis for the analyses that follow. Applying this framework to 31,143 VMRs, we classified high-confidence VMRs by local SNP-explained methylation variation and compared VMR classes across genomic annotation, chromatin state, repeat element overlap, trait heritability enrichment, local transcriptional associations, and available environmental and clinical metadata. This ancestry-aware analysis resolves components of brain methylation variation explained, or not explained, by nearby SNPs in an underrepresented population, distinguishing distal, repeat-rich genetically anchored methylation from gene-proximal, exposure-associated variation, with implications for interpreting epigenomic mechanisms relevant to neuropsychiatric and neurodegenerative risks (**Fig. 1**).

**Fig. 1.**
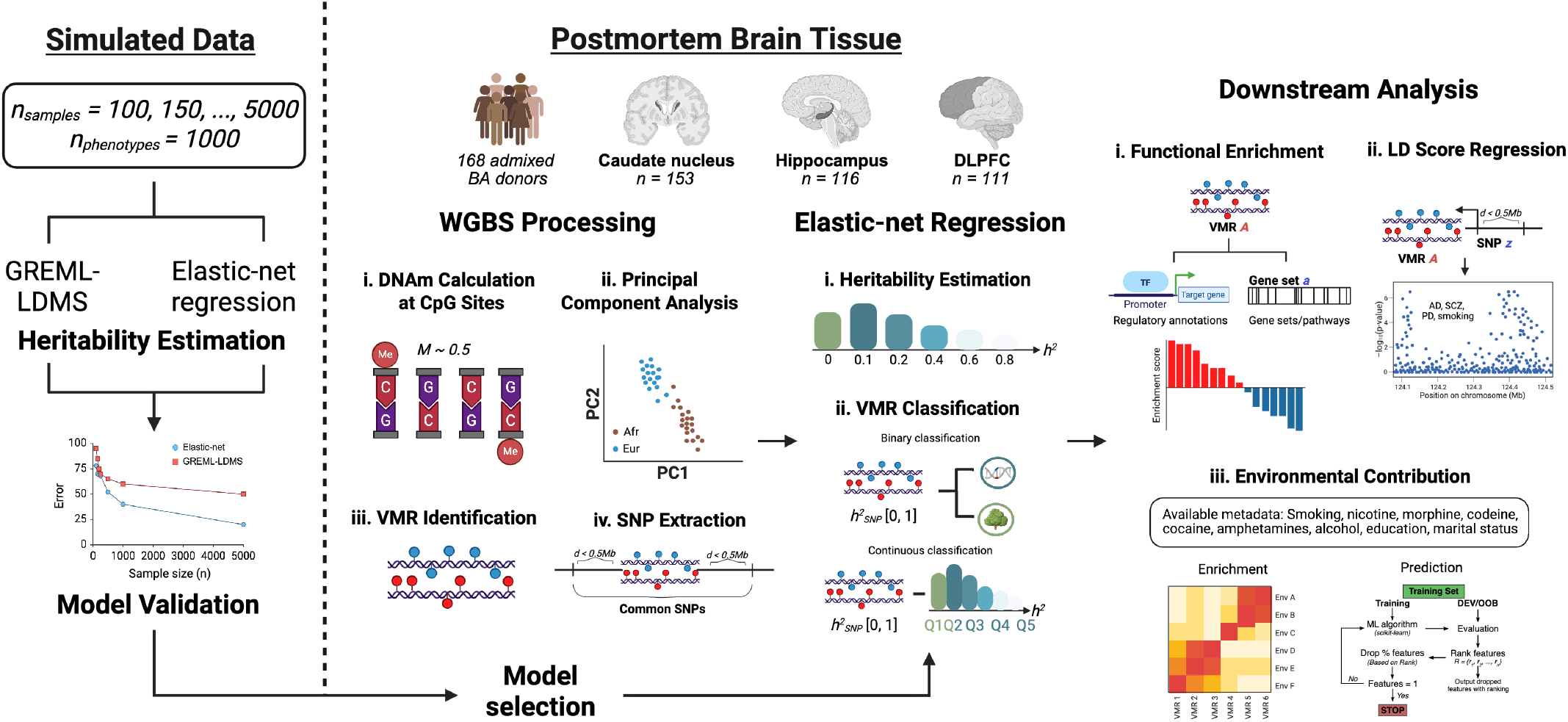
Overview of study design and analytical workflow. SNP-explained methylation variation 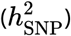 estimation was benchmarked using simulated DNA methylation (DNAm) phenotypes across seven sample sizes to compare elastic-net regression and linkage disequilibrium-stratified genomic restricted maximum likelihood (GREML-LDMS) performance. The benchmarking-informed framework was then applied to whole-genome bisulfite sequencing (WGBS) data from the BrainSEQ consortium postmortem hippocampus, caudate nucleus, and dorsolateral prefrontal cortex (DLPFC) tissue from Black American (BA) donors. Variably methylated regions (VMRs) were classified by local 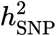 and prediction confidence (*r*^2^), then compared across genomic annotation, chromatin and repeat context, nearby gene expression and splicing associations, trait-level GWAS heritability, and available environmental and clinical metadata. Created with *BioRender.com*.

## Results

### A SNP-based modeling framework enables reliable VMR classification at postmortem brain sample sizes

To benchmark methods for estimating VMR-level 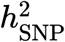, we compared GREML-LDMS with boosting elastic-net regression in simulations spanning sample sizes typical of large DNAm studies (*n* = 100 – 5000). We simulated 1,000 phenotypes, comprising 25% high-SNP (high local SNP-explained variation; 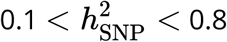) and 75% low-SNP (low SNP-explained variation; 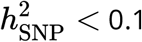) regions (**Fig. S1**). We classified each as high-SNP, low-SNP, or low-prediction on the basis of estimated 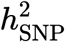 and model confidence (*r*^2^*threshold* = 0.3). Performance was assessed by concordance between simulated and estimated 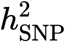, false-positive classification of low-SNP regions, and estimation bias across heritability bins.

At the sample sizes used in our downstream analyses (*n* = 150 – 250), elastic net returned stable estimates for all simulated phenotypes, with false-positive rates of 9.5–33.0% and power of 51.6– 73.9% (**Fig. 2a**). In contrast, GREML-LDMS frequently failed to produce stable estimates at *n* = 100 – 250 and remained underpowered at intermediate sample sizes (*n* = 500, 1000; **Fig. S2**), with limited correspondence between estimated and simulated 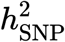 (**Fig. S3**). At intermediate sample sizes (*n* = 500, 1000), elastic net showed elevated false-positive rates (87.1% and 83.2%, respectively; **Fig. S2a**,**b**), reflecting increased misclassification of low-SNP regions as high-SNP outside the sample size range of interest. At *n* = 5000, the false-positive rate fell to 6.3%, but power also dropped to 34.4%, consistent with more stringent confidence thresholds excluding borderline signals. These benchmarks therefore motivated the use of boosting elastic net primarily as a classification and ranking framework for local SNP predictability, rather than as a calibrated estimator of absolute VMR-level heritability across all sample size regimes. Because region-level sample sizes in our postmortem cohort fell within the range in which elastic net yielded stable estimates, we used boosting elastic net for downstream 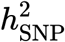 estimation and classification at DNAm sites.

**Fig. 2.**
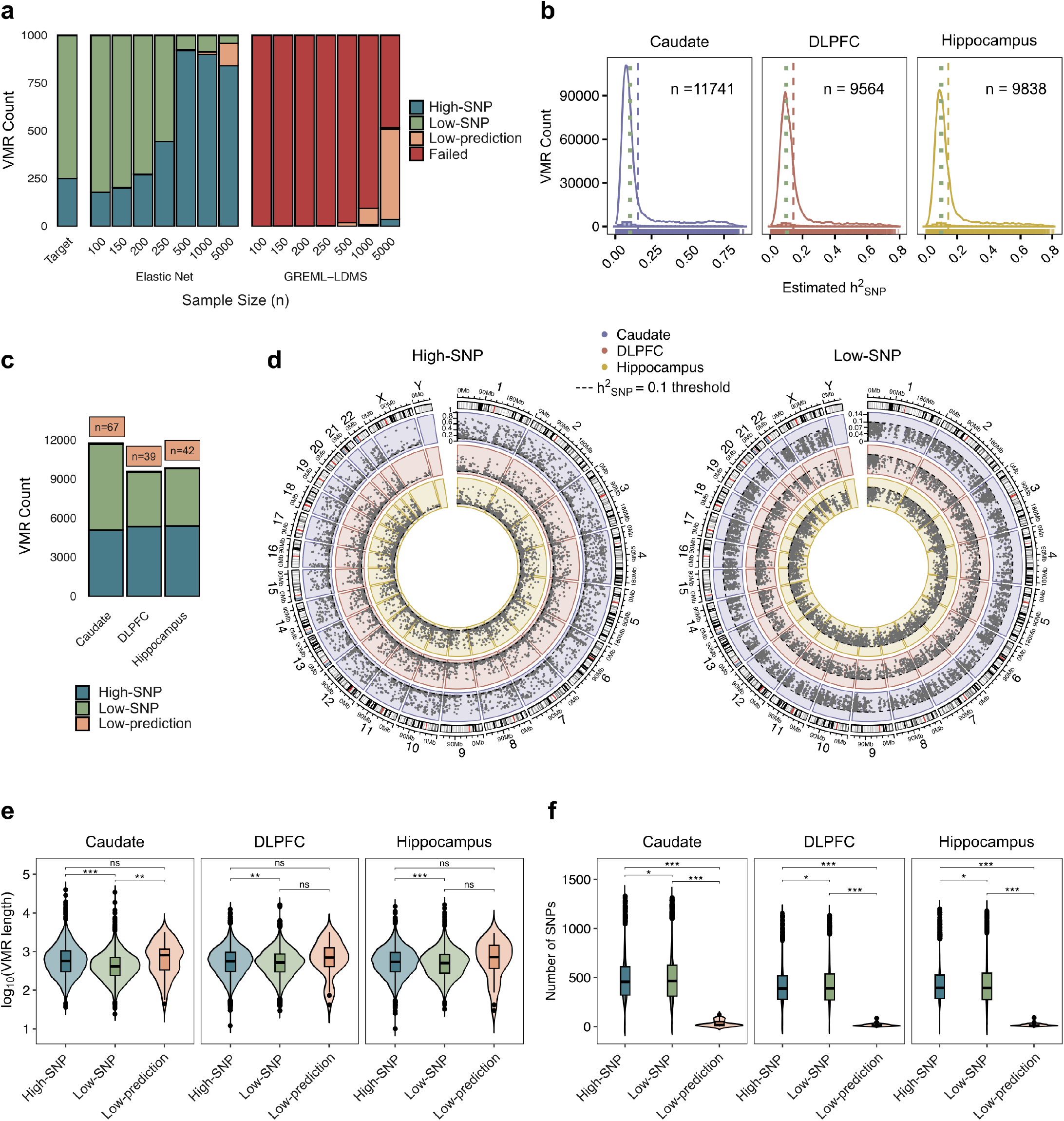
Brain VMRs partition into high-SNP, low-SNP, and low-prediction classes. **a**. Classification of simulated methylation phenotypes across sample sizes using elastic net and linkage disequilibrium-stratified genomic restricted maximum likelihood (GREML-LDMS). Colors indicate high local SNP-explained variation (high-SNP), low local SNP-explained variation (low-SNP), low-prediction, and failed classifications; the target bar shows the simulated class distribution. **b**. Distribution of estimated SNP-based heritability 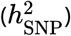 for retained variably methylated regions (VMRs) in the caudate, dorsolateral prefrontal cortex (DLPFC), and hippocampus. Dashed lines indicate the 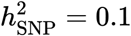 threshold. **c**. Counts of high-SNP, low-SNP, and low-prediction VMRs by brain region. **d**. Chromosomal distribution of high-SNP and low-SNP VMRs across brain regions. Points show regional 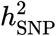 estimates. **e**., **f**. VMR length (**e**) and local SNP count (**f**) by SNP-explained class and brain region. Boxplots show medians and interquartile ranges; significance labels indicate adjusted pairwise comparisons (*P* < 0.05 (*), *P* < 0.01 (**), *P* < 0.001 (***), or ns [not significant]).

### Brain VMRs partition into reproducible SNP-explained classes

Using elastic net, we estimated 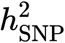 for DNAm in a BA discovery cohort from the BrainSEQ Consortium (*n* = 168 unique donors; **Table 1**) across three brain regions: DLPFC (*n* = 111), hippocampus (*n* = 116), and caudate nucleus (*n* = 153), after residualizing DNAm levels for genetic similarity, batch effects, age, sex, and schizophrenia case-control status (**Methods**). To enable genome-scale analysis, we quantified DNAm at VMRs rather than individual CpG sites. Across the genome, we identified 32,589 VMRs, including 10,372 in the DLPFC, 10,216 in the hippocampus, and 12,001 in the caudate nucleus (**Table S1**). After excluding VMRs without nearby SNPs within a ±500 kb window, removing additional loci lacking SNPs after clumping, and discarding 37 sites for which elastic net did not return an estimate, 31,143 VMRs remained for downstream analysis. The distribution of 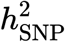 was similar across brain regions, spanning near-zero to ~0.9 and peaking at approximately 0.093 in the DLPFC, 0.089 in the hippocampus, and 0.075 in the caudate nucleus with respective mean values of 0.14, 0.15, and 0.16 (**Fig. 2b**).

**Table 1:** Discovery cohort. Black American (BA) sample characteristics for adult (age > 17) postmortem brain regions. Values are mean (SD) or n (%).

| Characteristic | Caudate (N = 153) | DLPFC (N = 111) | Hippocampus (N = 116) |
| --- | --- | --- | --- |
| Age | 50 (14) | 49 (14) | 50 (14) |
| Sex |  |  |  |
| Female | 54 (35%) | 46 (41%) | 48 (41%) |
| Male | 99 (65%) | 65 (59%) | 68 (59%) |
| **Dx** |  |  |  |
| *CTL* | 88 (58%) | 68 (61%) | 68 (59%) |
| *SCZ* | 65 (42%) | 43 (39%) | 48 (41%) |

We next classified VMRs using estimated 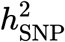 and associated prediction confidence (*r*^2^). VMRs with 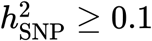 and *r*^2^ > 0.3 were defined as high-SNP, those with 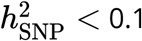 and *r*^2^ >0.3 as low-SNP, and those with *r*^2^ ≤ 0.3 as low-prediction. Under this framework, VMRs stratified into 15,802 high-SNP, 15,193 low-SNP, and 148 low-prediction sites across all brain regions (**Fig. 2d** and **Table S2**).

Regional distributions of these classes are summarized in **Fig. 2c** and **Table S3**. Classification was robust to a more stringent prediction threshold (*r*^2^ ≤ 0.75). Under this criterion, high-confidence high-SNP and low-SNP VMRs were largely concordant with the primary analysis, whereas most reclassified loci shifted to the low-prediction category, predominantly from the low-SNP class (**Fig. S7**).

To determine whether this classification simply reflected differences in local genetic input, we examined the number of SNPs extracted from each VMR and the length of the underlying intervals. Across all brain regions, low-SNP VMRs contained more SNPs on average than high-SNP or low-prediction VMRs (linear mixed-effects model, Bonferroni-corrected *P* < 0.05; **Fig. 2e,f**), arguing against insuffcient local SNP availability as an explanation for their classification. VMR length also differed across regions and SNP-explained classes (linear mixed-effects model, Bonferroni-corrected *P* < 2.2 × 10^−16^), but its correlation with estimated 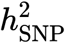 was weak in the caudate (*r* = 0.23, *FDR* = 5.94 × 10^−135^), DLPFC (*r* = 0.069, *FDR* = 1.56 × 10^−11^), and hippocampus (*r* 0.081, *FDR* = 1.05 ×^−15^) (**Fig. S6**). These findings indicate that the observed VMR classes are not explained by interval length or SNP density alone.

For replication in a multi-ancestry expanded cohort, we added 142 non-Hispanic white American (WA) donors (caudate *n* = 129,, DLPFC *n* = 55, hippocampus *n* = 60; **Table S4**) to the BA discovery set. VMRs retained the same broad class structure across brain regions, with 12,079 high-SNP, 5,034 low-SNP, and 182 low-prediction VMRs consistently classified within the BA and WA donor groups (**Table S5**). Cross-donor-group concordance of estimated 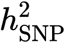 was consistently stronger for high-SNP than for low-SNP VMRs and varied markedly by region, with the caudate showing the highest correlation (Spearman’s *ρ* = 0.81) and DLPFC the lowest (*ρ* =0.26; hippocampus *ρ* = 0.46). Low-SNP VMRs were essentially uncorrelated across all regions (*ρ* = 0.03 –0.06; **Fig. S10**). Thus, VMRs classified as high-SNP showed greater cross-group consistency in estimated local SNP contribution—particularly in the caudate—whereas Low-SNP VMRs showed limited concordance, consistent with weaker shared local SNP effects.

### Low local SNP-explained VMRs show reduced cross-region concordance

We next examined the extent to which VMRs were shared across the DLPFC, caudate, and hippocampus (**Fig. 3a**). Across all VMRs, the caudate contained the largest number of region-specific loci (*n* = 6.639), whereas the DLPFC and hippocampus shared the largest common set (*n* = 6,936). Overlap between the caudate and either DLPFC (*n* = 4,423) or hippocampus (*n* = 4,499) was smaller, and only 3,395 VMRs were shared across all three regions. This overall pattern was preserved after stratification by SNP-explained class, with the caudate consistently showing the greatest region specificity and the DLPFC and hippocampus remaining the most concordant tissue pair. However, this also revealed increased regional specificity in the DLPFC and hippocampus in high-SNP (DLPFC, n = 2,549; hippocampus, n = 2,487) and low-SNP (DLPFC, n = 2,048; hippocampus, n = 2,238) VMR classes.

**Fig. 3.**
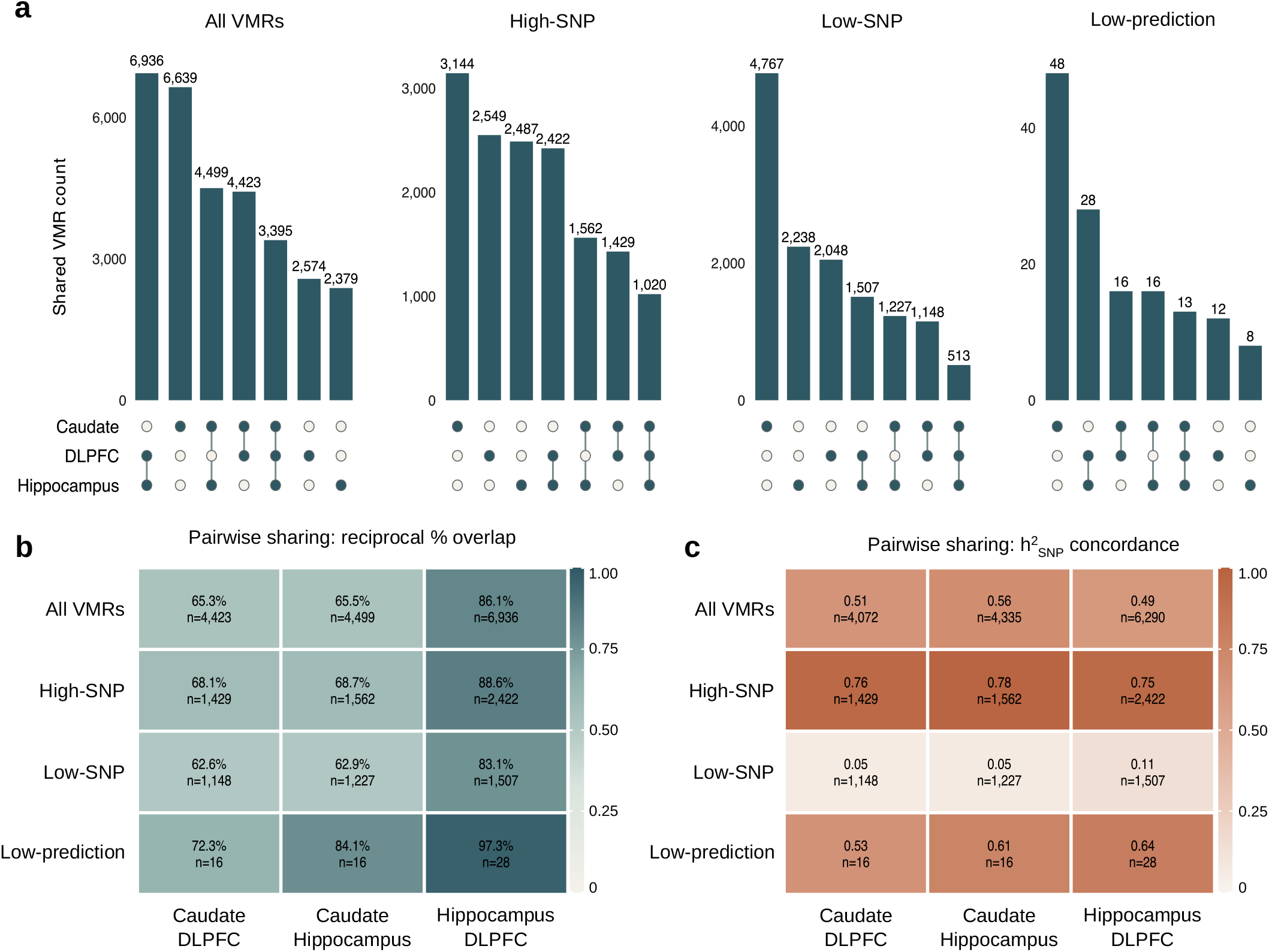
VMR sharing and local SNP-explained methylation variation concordance differ across brain regions. a. UpSet plots showing region-specific and shared variably methylated regions (VMRs) across the caudate, dorsolateral prefrontal cortex (DLPFC), and hippocampus for all VMRs and for each SNP-explained class. **b**. Pairwise reciprocal percentage overlap among VMRs shared between brain region pairs, stratified by SNP-explained class. Values show reciprocal overlap and the number of shared VMRs. **c**. Concordance of estimated SNP-based heritability 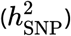 among shared VMRs across brain region pairs. Values show Spearman’s correlation coeffcient and the number of VMRs included in each comparison.

To quantify cross-region similarity, we calculated reciprocal percent overlap and Jaccard indices for each pairwise comparison, using fractional overlap criteria to account for differences in VMR lengths across regions. DLPFC and hippocampus showed the highest similarity across high-SNP, low-SNP, and low-prediction VMRs, whereas the caudate remained the most distinct region (**Fig. 3b**). Pairwise comparison of estimated 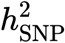 across overlapping VMRs showed moderate concordance overall (Spearman’s *ρ* = 0.28–0.40, *P* < 0.05) and substantially stronger correlations among high-SNP VMRs (Spearman’s *ρ* = 0.68–0.75,*P* < 0.05). By contrast, low-SNP VMRs showed no cross-region correlation in estimated 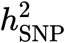 (Spearman’s *ρ* = −0.016 to 0.07, *P* > 0.05; **Fig. 3c**). Discordance rates and directional class shifts across region pairs are summarized in **Tables S6** and **S7**.

### Local SNP-explained variation increases across a distal intergenic regulatory gradient

Genomic annotation and pathway analyses identified distinct regulatory profiles for high-SNP and low-SNP VMRs across brain regions (**Fig. 4a**). High-SNP VMRs were consistently shifted away from gene-proximal regulatory features and toward intergenic regions, with the strongest signal observed in the caudate. In the caudate, high-SNP VMRs were enriched in intergenic (OR = 1.82, FDR< 10^−39^) and CpG-intergenic (OR = 1.74, FDR < 10^−48^) annotations and depleted from promoters, CpG islands, CpG shores, introns, exons, 5 UTRs, 3 UTRs, and enhancers (all FDR < 0.05). Hippocampal high-SNP VMRs showed a similar but weaker pattern, including enrichment in intergenic sequence (OR = 1.32, FDR < 10^−7^) and depletion from several gene-linked features. In the DLPFC, annotation signals were more limited, with significant depletion restricted to 5 UTRs (OR = 0.77, FDR = 0.030).

**Fig. 4.**
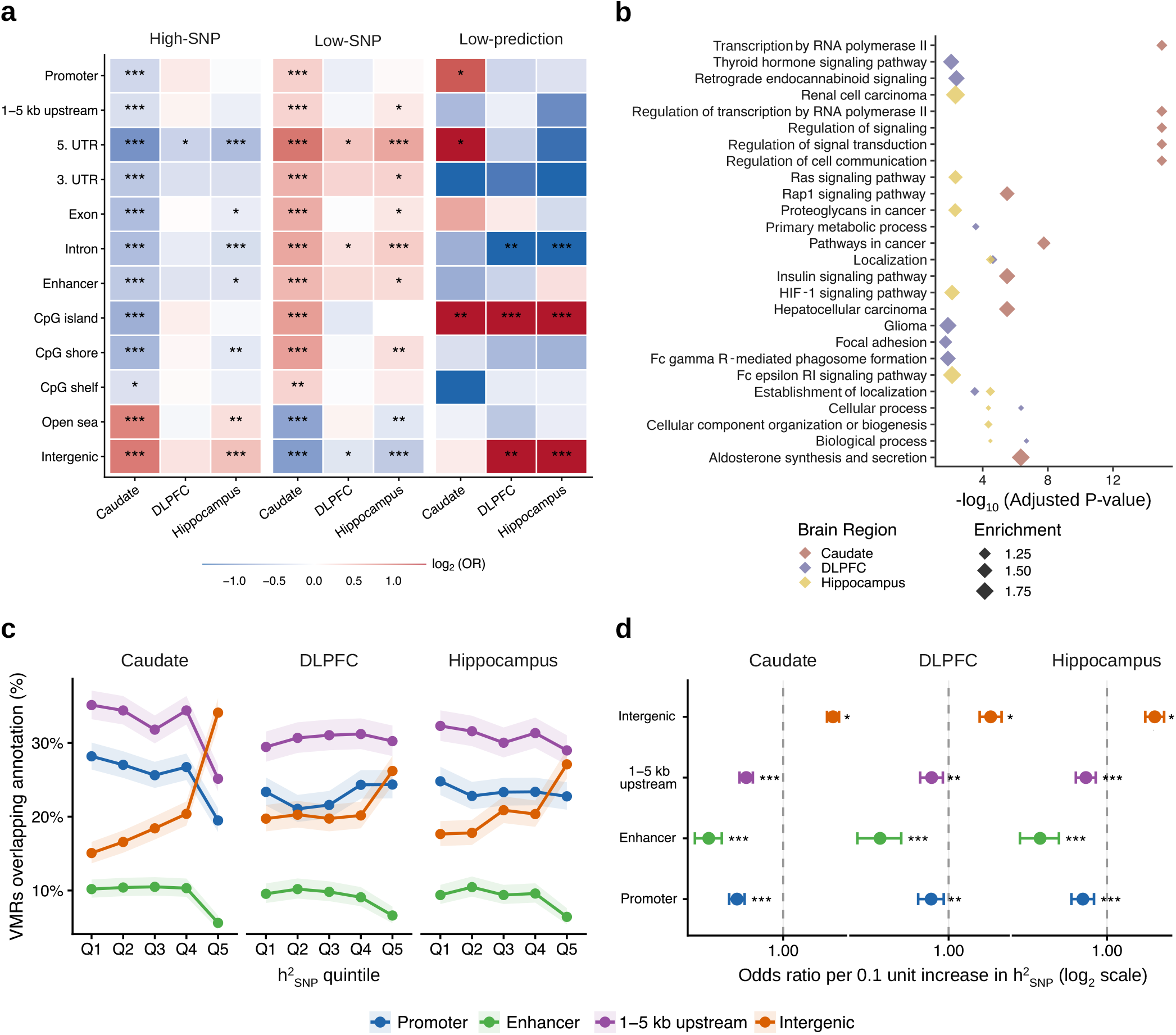
High-SNP VMRs are shifted toward distal intergenic annotations. **a**. Enrichment and depletion of variably methylated region (VMR) SNP-explained classes across genomic and regulatory annotations in the caudate, dorsolateral prefrontal cortex (DLPFC), and hippocampus. Colors show log2-transformed odds ratios (ORs) from Fisher’s exact tests. **b**. Enriched gene set and pathway terms associated with VMR SNP-explained classes across brain regions, ranked by FDR-adjusted significance. **c**. Percentage of VMRs overlapping intergenic, 1–5 kb upstream, enhancer, and promoter annotations across 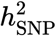 quintiles. Shaded areas indicate 95% confidence intervals (CIs). **d**. Logistic regression estimates for annotation overlap per 0.1-unit increase in 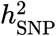, adjusted for VMR length and local SNP density. Points show log2-transformed ORs; error bars show 95% CIs. FDR-adjusted p-values are indicated by *P* < 0.05 (∗), *P* < 0.01 (∗∗), or *P* < 0.001 (∗∗∗)

Low-SNP VMRs showed a broadly reciprocal profile (**Fig. 4a**). In the caudate, low-SNP VMRs were enriched in promoters, CpG islands, CpG shores, introns, exons, 5 UTRs, and enhancers (all FDR 0.05) and were depleted from intergenic sequences (OR = 0.55, FDR < 10^−39^). Hippocampal low-SNP VMRs were also depleted from intergenic regions (OR = 0.73, FDR < 10^−9^) and enriched in intronic and 5 UTRs. In the DLPFC, low-SNP VMRs showed weaker and less consistent annotation differences. Low-prediction VMRs displayed sparse and heterogeneous annotation signals; although, CpG island depletion was observed across brain regions.

Pathway analysis further separated the major VMR classes (**Fig. 4b** and **Fig. S11**). High-SNP VMRs were enriched for broad metabolic and cellular processes and for Huntington’s disease-associated gene sets, consistent with the prominent caudate signal in this class. Low-SNP VMRs were enriched for neurological pathways, including spinocerebellar ataxia and retrograde endocannabinoid signaling, whereas low-prediction VMRs showed enrichment for neurodegeneration- and neurotrophin-related pathways. Full pathway results are provided in **Table S8**.

We next tested whether regulatory annotation varied continuously across the 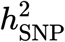 spectrum. Restricting to high-confidence VMRs with cross-validated *r*^2^ >0.3, we modeled annotation membership as a function of 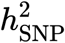 using logistic regression adjusted for VMR length and SNP density. Intergenic overlap increased monotonically with 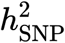 in all three brain regions with ORs per 0.1-unit increase in 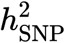 of 1.23 in the caudate (95% CI, 1.20–1.26), 1.15 in the DLPFC (95% CI, 1.11–1.19), and 1.19 in the hippocampus (95% CI, 1.15–1.23; all FDR < 10^−13^; **Fig. 4c**). Conversely, overlap with promoter, enhancer, and upstream (1–5 kb) regions decreased with increasing 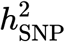 (all FDR 0.05; **Fig. 4d**). These associations were supported by likelihood-ratio tests comparing natural cubic spline models with covariate-only nulls and were robust to inclusion of low-prediction VMRs in a sensitivity analysis (**Fig. S13**).

The intergenic gradient was also observed in the multi-ancestry matched cohort. Across both BA and WA donor groups, intergenic enrichment increased with 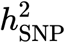 and reciprocal depletion of promoter, enhancer, and 1–5 kb upstream annotations (**Fig. S15**; **Table S9**). Together, these results indicate that increasing 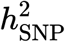 tracks with a shift away from gene-proximal regulatory annotations and toward distal intergenic sequences.

### Highly local SNP-explained intergenic VMRs mark repeat-rich, repressive chromatin

Because high-SNP VMRs were preferentially located in intergenic sequences, we next asked whether these loci overlapped brain cell-type-specific open chromatin. Using PsychENCODE BrainScope single-nucleus Assay for Transposase-Accessible Chromatin (ATAC)-seq peaks from seven brain cell types [22], we found that high-SNP intergenic VMRs were consistently depleted from open chromatin relative to low-SNP intergenic VMRs across cell types and brain regions (Fisher’s exact test, all FDR < 0.05; **Fig. 5a**). In the pooled analysis, 57% of high-SNP intergenic VMRs overlapped the union ATAC peak set, compared with 76% of low-SNP intergenic VMRs (OR = 2.36, FDR <10^−52^, where OR >1 indicates enrichment outside ATAC-defined open chromatin). This depletion was most pronounced in glial cell types, including oligodendrocyte progenitor cells (OR = 2.06, FDR <10^−42^), oligodendrocytes (OR = 2.03, FDR <10^−39^), and astrocytes (OR = 1.89, FDR <10^−32^), and was weaker in microglia (OR = 1.44, FDR <10^−9^). Across brain regions, the signal remained directionally consistent but was strongest in the caudate (union OR = 4.25, FDR <10^−56^) and more modest in the DLPFC and hippocampus (union OR = 1.72 and 1.67, respectively).

**Fig. 5.**
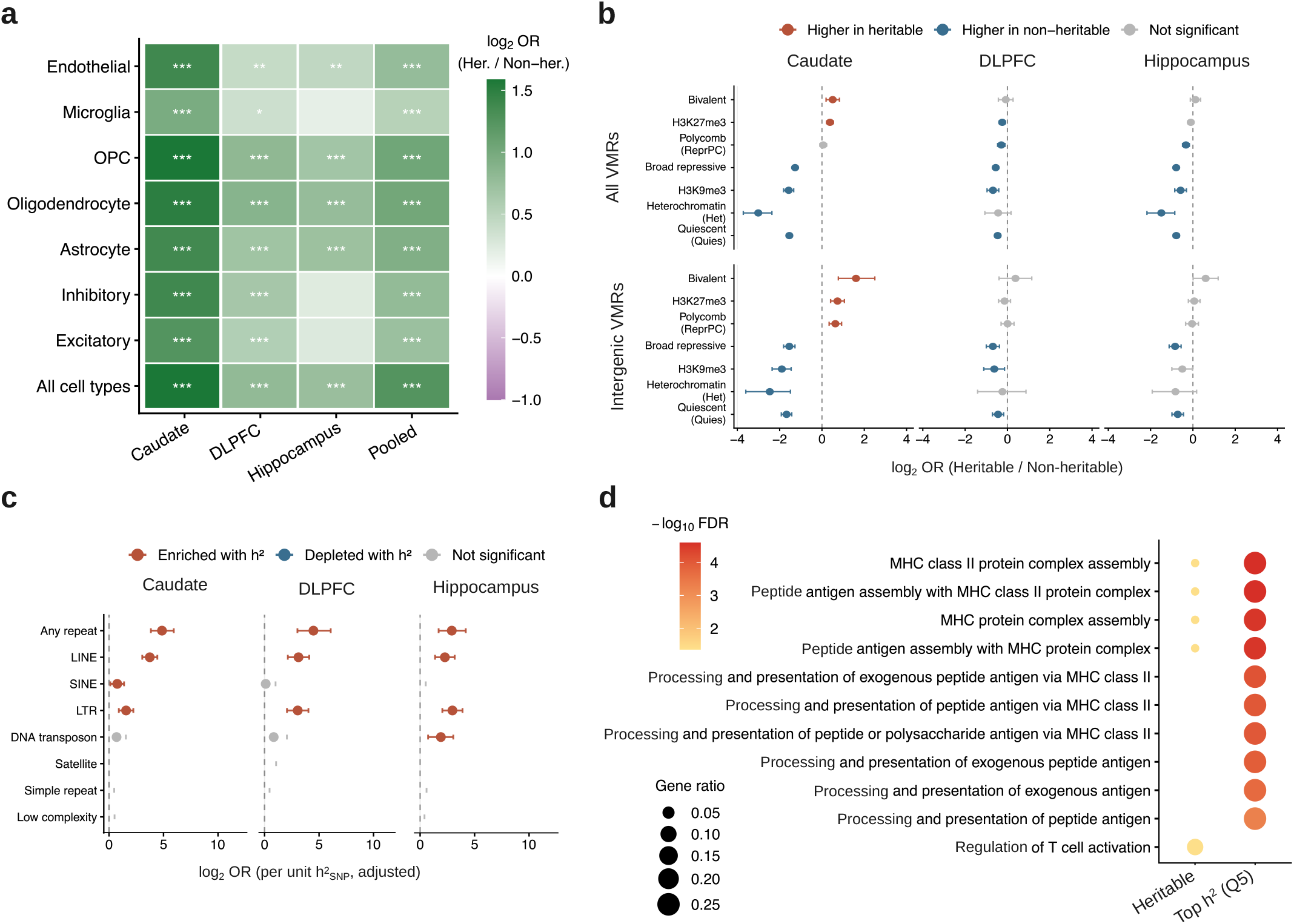
Highly local SNP-explained intergenic VMRs are enriched in repressive and repeat-rich chromatin. **a**. Open chromatin enrichment or depletion for high-SNP versus low-SNP intergenic variably methylated regions (VMRs) across brain cell types and regions. Colors show log2-transformed odds ratios (ORs). **b**. Enrichment of high-SNP versus low-SNP VMRs in repressive chromatin features, shown for all VMRs and intergenic VMRs across the caudate, dorsolateral prefrontal cortex (DLPFC), and hippocampus. **c**. Association between continuous SNP-based heritability 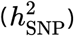 and repetitive element overlap among intergenic VMRs, adjusted for VMR length and local SNP density. **d**. Gene set enrichment among genes linked to high-SNP intergenic VMRs and top-quintile (Q5) high-SNP intergenic VMRs. Dot size and color indicate enrichment strength and FDR-adjusted significance, respectively.

This pattern extended across the continuous 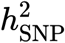 spectrum (**Fig. S13**). Among intergenic VMRs, 68– 77% of VMRs in the first four 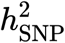 quintiles overlapped the union ATAC peak set, compared with only 32% of VMRs in the top quintile (Q5; median 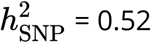). Thus, Q5 intergenic VMRs were markedly depleted from open chromatin relative to Q1–Q4 VMRs (OR = 0.17 for ATAC overlap, FDR <10^−157^). Logistic regression confirmed a negative continuous association between 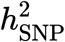 and open chromatin overlap across cell types and brain regions (all FDR <10^−5^).

To define the chromatin context of these loci more broadly, we intersected VMR coordinates with repressive histone mark peaks and ChromHMM chromatin state annotations from the NIH Roadmap Epigenomics Consortium [23,24], using tissue-matched brain epigenomes for the caudate, DLPFC, and hippocampus (**Fig. 5b**). Across all brain regions, overlap with quiescent chromatin showed the strongest positive association with 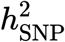, after adjustment for VMR length and SNP density. Per-unit increases in 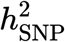 were associated with substantially greater odds of quiescent state overlap in the caudate (OR = 69.8, 95% CI, 53.1–92.4; FDR = 3.9 × 10^−198^), hippocampus (OR = 42.5, 95% CI, 30.1– 60.2; FDR = 4.9 × 10^−100^), and DLPFC (OR = 21.4, 95% CI, 14.9–30.8; FDR = 7.7 × 10^−62^).

Patterns of repressive histone marks differed between the broader high-SNP intergenic class and the highest-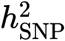 subset. In the caudate, high-SNP intergenic VMRs were enriched relative to low-SNP intergenic VMRs in H3K27me3-marked and Polycomb-repressed chromatin (H3K27me3: OR = 1.66, FDR = 1.3 × 10^−5^; Polycomb state: OR = 1.55, FDR = 3.9 × 10^−5^), whereas H3K9me3-marked chromatin was depleted in the high-SNP class overall. In contrast, Q5 intergenic VMRs were enriched for H3K9me3-marked chromatin relative to Q1–Q4 VMRs across brain regions (caudate OR = 6.32, FDR = 1.4 × 10^−66^; DLPFC OR = 2.08; hippocampus OR = 3.35) and depleted for H3K27me3-marked chromatin in the caudate (OR = 0.26, FDR = 3.9 × 10^−14^). These results indicate that the most highly high-SNP intergenic VMRs are concentrated in quiescent and repressive chromatin annotations.

This chromatin architecture coincided with strong enrichment for LINE elements, particularly the L1 subfamily (**Fig. 5c**). Compared with Q1–Q4 intergenic VMRs, Q5 intergenic VMRs showed substantially higher LINE overlap in the caudate (OR = 9.49, FDR = 5.68 × 10^−88^), DLPFC (OR = 3.10, FDR =2.19 × 10^−16^), and hippocampus (OR = 2.81, FDR = 7.39 × 10^−14^). In absolute terms, 65.4%, 43.6%, and 41.4% of Q5 intergenic VMRs overlapped L1 elements in the caudate, DLPFC, and hippocampus, respectively. Continuous models also showed a positive monotonic association between 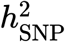 and LINE overlap across all brain regions (**Fig. S13**).

These chromatin signatures were recapitulated in the multi-ancestry matched cohort. In both BA and WA donor groups, quiescent ChromHMM state remained the strongest positive association with 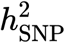, low-SNP intergenic VMRs showed greater open chromatin overlap than high-SNP intergenic VMRs, and the top 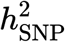 quintile remained strongly depleted from ATAC peaks relative to Q1–Q4 VMRs. LINE/L1 enrichment in the top 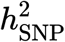 quintile was directionally consistent across regions, with L1 overlap observed for 72.2%, 34.8%, and 47.2% of Q5 intergenic VMRs in the caudate, DLPFC, and hippocampus, respectively (**Fig. S16**; **Fig. S15**; **Table S10**).

Finally, we asked whether the minority of high-SNP intergenic VMRs that overlapped putative regulatory elements could be linked to candidate target genes. Of 3,532 pooled high-SNP intergenic VMRs, 101 (~3%) overlapped Activity-by-Contact (ABC)-defined enhancers, yielding 179 VMR–gene links to 119 unique target genes. The corresponding low-SNP set showed similar enhancer overlap (102 VMRs, 263 links, 158 genes), with no difference in ABC scores (Wilcoxon *P*= 0.42). Genes linked to high-SNP intergenic VMRs were enriched for MHC class II antigen presentation terms, including MHC class II protein–complex assembly (*HLA-DQA1/DQB1/DRB1*; FDR = 0.043; **Fig. 5d**) and regulation of T cell activation (FDR = 0.043). This signal was stronger among Q5 VMR-linked genes, although the number of mapped target genes was small. By contrast, genes linked to low-SNP intergenic VMRs showed no significant pathway enrichment. Together, these results indicate that highly local SNP-explained brain VMRs are concentrated in repressive, repeat-rich intergenic chromatin, with a small candidate regulatory subset linked to immune-related target genes.

### Distal high-SNP VMRs show nearby transcriptional and splicing associations

The genomic annotation and chromatin analyses indicated that high-SNP VMRs are enriched in distal, repressive, repeat-rich intergenic sequence, whereas low-SNP VMRs are preferentially located near gene-proximal regulatory elements. We therefore asked whether this spatial partitioning was accompanied by differences in local molecular associations. We tested pairwise associations between VMR methylation levels and nearby transcriptional features using two complementary strategies: gene expression, assessed through ABC enhancer–promoter links and a 250 kb nearest gene window, and alternative splicing, quantified as percent-spliced-in (PSI) for events within 250 kb.

For gene expression, the number of testable VMR–gene pairs under the ABC strategy was limited, consistent with the sparse overlap between intergenic VMRs and ABC-defined enhancers described above. In the caudate and hippocampus, high-SNP VMRs had numerically higher ABC-linked discovery rates than low-SNP VMRs (caudate: 5.9% high-SNP [2 of 34] vs. 2.0% low-SNP [1 of 50]; hippocampus: 2.6% high-SNP [1 of 38] vs. 0% low-SNP [0 of 25]; FDR <0.05). No significant ABC-linked associations were detected in the DLPFC. Using the nearest gene strategy, which substantially expanded the testable pair set, high-SNP VMRs showed higher discovery rates than low-SNP VMRs across brain regions, considering significant VMR–gene associations at FDR <0.05 (caudate: 4.1% [170 of 4,111] vs. 2.3% [133 of 5,794]; DLPFC: 1.7% [76 of 4,471] vs. 0.42% [15 of 3,583]; hippocampus: 1.7% [75 of 4,469] vs. 0.63% [24 of 3,816]).

Methylation–splicing associations were also more frequently detected among high-SNP VMRs. In the caudate, 2.3% of tested high-SNP VMRs were associated with at least one PSI event at FDR <0.10 (114 of 4,938), compared with 1.0% of low-SNP VMRs (63 of 6,579; **Fig. S18**). This pattern was directionally consistent, although weaker, in the DLPFC (high-SNP: 0.59% [31 of 5,291] vs. low-SNP: 0.41% [17 of 4,158]) and hippocampus (0.54% [29 of 5,340] vs. 0.46% [20 of 4,376]). Median peak effect sizes were larger for methylation–splicing associations than for methylation–expression associations (median |*β*|_max_ ≈ 2.8 versus ≈0.5 on their respective normalized scales). These results suggest that high-SNP VMRs show detectable local coupling to both gene expression and alternative splicing, with the strongest splicing enrichment observed in the caudate.

A complementary genomic distance analysis confirmed that high-SNP and low-SNP VMRs differ in their spatial relationship to transcribed features across brain regions. Low-SNP VMRs were located closer to the nearest gene transcription start site (TSS) than high-SNP VMRs across brain regions (Wilcoxon, *P* < 0.05) with the caudate showing the strongest associations. Moreover, low-SNP VMRs showed the strongest enrichment for gene-proximal localization in the caudate (OR = 2.28; *P* = 4.86 × 10^−22^), with smaller but significant effects in the hippocampus (OR = 1.33; FDR = 0.003) and DLPFC (OR = 1.24; FDR = 0.030) after adjustment for VMR length and local SNP density. Distance to the nearest TSS increased monotonically across 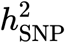 quintiles in all three regions (*P* < 1.1 × 10^−5^), consistent with a continuous relationship between local 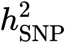and gene distance rather than a discrete threshold effect. A similar pattern was observed for proximity to tested splicing events. Low-SNP VMRs were more likely than high-SNP VMRs to be proximal to splicing events in the caudate and hippocampus (FDR < 0.005), with a directionally consistent but non-significant effect in the DLPFC (FDR = 0.061). Distance to the nearest PSI event also increased across 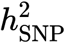 quintiles in all brain regions (*P* < 2.2 × 10^−4^; **Fig. S18**).

These local regulatory patterns were partially recapitulated in the multi-ancestry matched cohort. Replication was strongest in the caudate, where low-SNP VMRs remained closer to genes and splicing events than high-SNP VMRs (OR = 2.67, FDR ≈ 8 × 10^−20^ and OR = 1.61, FDR, ≈ 4 × 10^−18^, respectively). Distance increased across 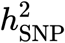 quintiles (Kruskal *P* = 8.4 × 10^−58^ and *P* = 1.2 × 10^−59^, respectively), and high-SNP VMRs retained higher nearest gene and PSI discovery rates than low-SNP VMRs (**Fig. S19**; **Table S11**). Hippocampal proximity contrasts remained directionally consistent but attenuated, whereas DLPFC proximity contrasts were not significant in the matched set.

Together, these analyses demonstrate that low-SNP VMRs are preferentially located near genes and tested splice events, whereas high-SNP VMRs are located farther from transcribed features, consistent with their enrichment in distal repressive chromatin. Despite this greater genomic distance, high-SNP VMRs showed higher discovery rates for nearby gene expression and splicing associations. These patterns indicate that VMR SNP-explained classes differ not only in chromatin context and genomic position but also in the transcriptional features associated with local methylation variation.

### VMR SNP-explained classes show context-dependent metadata associations

To test whether VMR SNP-explained classes capture methylation variation associated with environmental, toxicological, sociodemographic, or clinical metadata, we first associated VMR methylation levels with available metadata variables in the BA discovery cohort. We then tested whether metadata-associated VMRs were enriched for high-SNP or low-SNP classes within each brain region. Variable definitions and sample coverage are provided in **Table S12**.

Metadata enrichments were strongly region- and class-dependent (**Fig. 6a,b**; **Fig. S20**; **Table S14**). In the caudate, high-SNP VMRs were enriched among VMRs associated with toxicology-defined opioid exposure (codeine OR = 1.62, FDR < 6×10^-5^; morphine OR = 1.46, FDR = 0.003), cocaine exposure (OR = 1.51, FDR = 0.003), amphetamine exposure (OR = 1.27, FDR = 0.024), and marital status but depleted among nicotine-associated VMRs (OR = 0.72, FDR < 2×10^-5^). In the DLPFC, high-SNP VMRs were enriched for ethanol-, amphetamine-, and codeine-associated VMRs, whereas marital-status-associated VMRs were enriched among low-SNP VMRs. In the hippocampus, high-SNP VMRs were enriched for educational attainment, morphine exposure, and cocaine exposure but depleted among nicotine- and amphetamine-associated VMRs. Thus, high-SNP and low-SNP VMRs did not map onto a single metadata axis; instead, their enrichments depended on brain region, metadata domain, and VMR class.

**Fig. 6.**
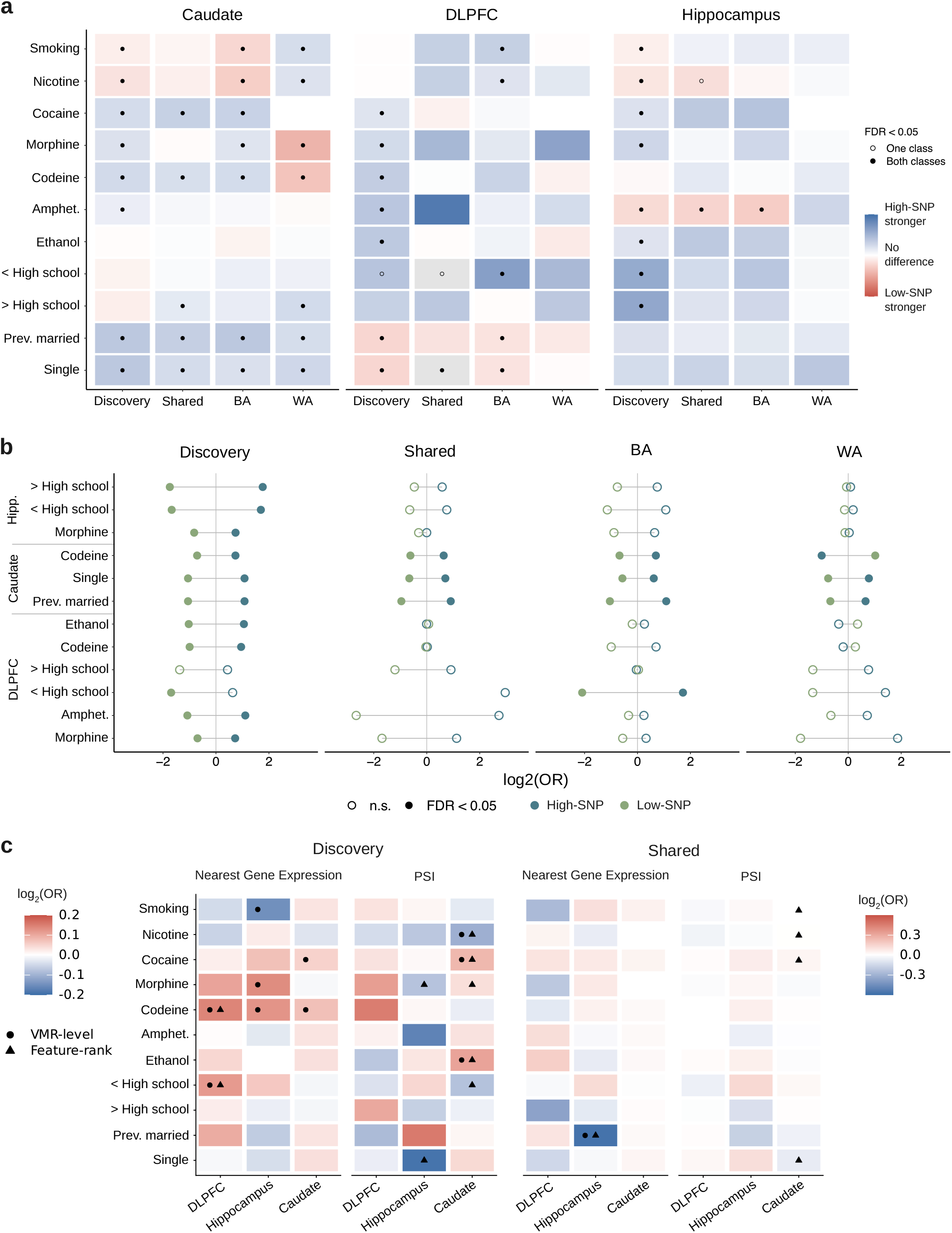
Environmental and clinical metadata associations vary by VMR SNP-explained class and brain region. **a**. Enrichment contrasts for metadata-associated variably methylated regions (VMRs) across discovery and shared analyses in the caudate, dorsolateral prefrontal cortex (DLPFC), and hippocampus. Colors indicate log2-transformed odds-ratio (OR) differences between high-SNP and low-SNP VMR enrichment; symbols denote FDR significance. **b**. Paired enrichment estimates for selected metadata variables comparing high-SNP and low-SNP VMR classes across discovery and shared analyses. Filled points indicate FDR-significant enrichments. **c**. Convergence between metadata-associated signals and VMR-linked transcriptional features. Heatmaps summarize associations for nearest gene expression and percent-spliced-in (PSI)-linked features across discovery and shared analyses; symbols distinguish VMR-level and feature-rank evidence.

Stratified analyses in the multi-ancestry matched cohort supported both shared and donor-group-dependent metadata enrichment patterns. In the caudate, marital-status- and educational-attainment-associated VMRs showed directionally consistent enrichment among high-SNP VMRs in both BA and WA donors (**Fig. S21-S23**\*; **Table S15**). By contrast, substance-use-associated signals were less consistent across donor groups. Codeine- and morphine-associated enrichments observed among BA high-SNP VMRs were attenuated or reversed in WA donors, whereas nicotine-associated patterns showed the opposite direction. These differences reduced the corresponding signals in combined analyses, indicating that pooled models can obscure donor-group-dependent enrichment patterns. Because these contrasts may reflect differences in exposure distributions, co-exposures, sample composition, or genetic architecture, we interpret them as consistent with context dependence rather than definitive ancestry-specific effects.

We next asked whether multivariate VMR signatures could recover metadata-associated structure across cohorts. Dynamic recursive feature elimination (dRFE) identified educational-attainment-associated high-SNP VMR signatures with modest cross-cohort performance in the hippocampus and caudate (**Table S18**). Absolute AUC values were generally low (0.50–0.67), consistent with the expected limited power of epigenome-wide methylation signatures to classify complex psychosocial exposures from cross-sectional postmortem data. Educational attainment was the only metadata variable with consistent predictive performance across discovery and expanded-cohort analyses (AUC ≥ 0.58 in both cohorts for caudate and hippocampus high-SNP VMRs). Other variables, including marital status, smoking, and nicotine exposure, exceeded the pre-specified threshold in discovery-only analyses but did not show consistent performance in the expanded cohort. These results suggest that educational-attainment-associated methylation structure was the most reproducible metadata signal, whereas substance-use-associated signatures were more sensitive to cohort composition.

Finally, we tested whether transcriptional features linked to high-SNP or low-SNP VMRs converged on the same environmental and clinical metadata. These analyses provided weaker and more context-specific associations than the VMR-level enrichment tests (**Fig. 6c**). Rank-based and VMR-level approaches identified recurrent signals involving substance use and psychosocial variables, including opioid, cocaine, ethanol, and nicotine exposure; smoking; educational attainment; and marital status with several associations reaching FDR < 0.10. However, thresholded overlap enrichment was limited. Thus, although SNP-explained classes showed distinct molecular coupling profiles, their linked transcriptional features converged only selectively with the environmental and clinical metadata available in this cohort.

Together, these results indicate that metadata-associated methylation variation is structured by SNP-explained class, brain region, and donor group. Educational attainment showed the clearest cross-cohort consistency, whereas substance-use-associated signals were more context-dependent, and convergence at the transcriptional-feature level was selective rather than broadly enriched.

### VMR SNP-explained classes show limited trait-level genetic overlap

The heterogeneous metadata associations described above motivated an orthogonal genetic contrast using stratified LD score regression (S-LDSC), which tests whether VMR annotations are enriched or depleted for trait-level GWAS heritability rather than individual-level metadata in this postmortem cohort. We tested VMR annotations stratified by binary SNP-explained class and by donor-group-defined 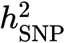 quintiles for enrichment or depletion of genome-wide association study (GWAS) heritability for smoking, AD, Parkinson’s disease (PD), and schizophrenia. Smoking provided a trait-level genetic contrast for addiction-related metadata, whereas AD, PD, and schizophrenia tested overlap with neurological and psychiatric trait architectures.

Binary SNP-explained classes showed no significant S-LDSC enrichment or depletion. In exploratory quintile analyses, however, several nominal signals were observed across brain regions and cohorts (**Fig. S24**; **Table S19**). In the BA discovery cohort, Q1 VMRs showed nominal depletion for PD (*z* = −2.70, *P* = 0.007) and schizophrenia (*z* = −2.52, *P* = 0.01) in the DLPFC. Similarly, in the multi-ancestry cohort, BA-defined Q3 VMRs showed nominal depletion for PD in the caudate (*z* = −2.93, *P* =0.003). Nominal enrichment signals included smoking in both Q1 (*z* =1.99, *P* =0.05) and Q2 (*z* =2.04, *P* = 0.04) DLPFC VMRs. Additionally, WA-defined Q4 VMRs showed nominal schizophrenia enrichment in the caudate (*z* =2.15, *P* =0.03), while Q5 VMRs showed nominal smoking depletion in the hippocampus (*z* = −2.46, *P* =0.01).

Together, these analyses indicate that VMR SNP-explained classes show limited and context-dependent overlap with trait-level GWAS heritability. Exploratory quintile analyses identified hypothesis-generating enrichment and depletion signals for smoking and neuropsychiatric or neurodegenerative traits, but these patterns were not observed as robust associations across binary SNP-explained classes and did not survive multiple-testing correction. These results suggest that local SNP-explained methylation variation does not map broadly onto the GWAS architectures of the tested traits, while highlighting specific brain-region and donor-group contexts for future investigation.

## Discussion

Brain DNAm variability in this underrepresented BA cohort is organized along a local SNP-explained methylation variation axis, ranging from distal, genetically anchored chromatin domains to gene-proximal variation less explained by nearby SNPs. Partitioning the BrainSEQ WGBS methylome by the degree of methylation variation explained by local SNP effects separated VMRs into high-SNP, low-SNP, and low-prediction classes that differed systematically in cross-region concordance, regulatory annotation, chromatin state, transcriptional coupling, and metadata association. These results support a model in which local genetic architecture does not act uniformly across the brain methylome, but instead marks distinct regulatory and chromatin compartments with different degrees of regional conservation, gene-proximal activity, and environmental or clinical association. By establishing that this partitioning can be performed robustly at postmortem brain sample sizes, the benchmarking analysis provided a principled foundation for resolving biologically distinct classes of SNP-associated methylation variation across the brain methylome.

An important feature of this framework is that local SNP-explained VMR behaves as a continuous spectrum rather than a binary property. The high-SNP, low-SNP, and low-prediction classes are pragmatic analytical categories used for enrichment testing, not discrete biological states. Consistent with this interpretation, functional annotations shifted progressively across the 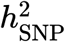 distribution: increasing heritability was associated with greater intergenic overlap, reduced promoter and enhancer overlap, lower open-chromatin overlap, and greater repeat element enrichment. Thus, VMRs near the 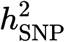 threshold should not be interpreted as categorically distinct from those just below it. Instead, the class-level differences observed here reflect aggregate trends across thousands of regions, with the strongest biological contrasts emerging at the upper end of the heritability spectrum.

The functional architecture of high-SNP VMRs points to genetically anchored distal methylation variation rather than canonical gene-proximal regulatory activity. Across brain regions, higher-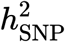 VMRs were progressively enriched in intergenic sequences and depleted from promoters, enhancers, CpG islands, and other gene-linked annotations. High-SNP VMRs also showed stronger cross-region concordance of estimated 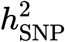 than low-SNP VMRs, indicating that the genetically anchored component of methylation variability is more reproducible across brain regions than the component not explained by local SNPs. This shared architecture was strongest between the DLPFC and hippocampus and weaker for comparisons involving the caudate, consistent with broader molecular and cellular distinctions between cortical/hippocampal and striatal tissue [25,26]. S-LDSC further suggested that quintile-based VMR annotations may show nominal, context-dependent overlap with trait-level GWAS heritability, but these exploratory signals did not survive multiple-testing correction and should be interpreted cautiously.

Integrating chromatin state, open chromatin, and repeat element annotations revealed that the highest-heritability intergenic VMRs are concentrated in closed, repeat-rich chromatin. High-SNP intergenic VMRs were depleted from brain cell-type-specific ATAC-seq peaks [22], and this depletion was strongest in the top heritability quintile, where only a minority of VMRs overlapped open chromatin. These loci were instead enriched for quiescent and constitutively repressive chromatin states, H3K9me3, and LINE/L1 elements. This pattern suggests that the strongest local genetic contribution to brain methylation occurs at distal loci embedded in repressive chromatin rather than at accessible enhancers or promoters [4,5,6,27]. Although the cohort included both neurotypical controls and donors with schizophrenia, schizophrenia case-control status was modeled explicitly: methylation was residualized for diagnosis prior to VMR-level 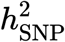 estimation (**Methods**). This adjustment makes it unlikely that the repeat-associated heterochromatin enrichment of high-SNP VMRs merely reflects uncorrected case-control methylation contrast. One plausible mechanism is that local genetic variation influences methylation through repeat-silencing pathways, including *KZFP/TRIM28/SETDB1*-mediated H3K9me3 deposition and *DNMT3A/3B* recruitment at transposable elements [28,29,30,31]. A complementary mechanism may involve *DNMT3A* recruitment to intergenic H3K36me2-marked regions through its PWWP domain [32]. Although these mechanisms were not directly tested here, they provide a biologically coherent model for why high-SNP methylation variation would concentrate in closed, LINE-rich intergenic sequences rather than in active regulatory elements. Taken together, these findings suggest that the most genetically anchored component of brain methylation variability is concentrated in genomic compartments involved in long-term epigenomic stability, heterochromatin maintenance, and repeat silencing.

This localization has implications for neuronal aging and neurodegeneration. High local SNP-explained VMRs concentrate in the distal, H3K9me3-marked, LINE/L1-rich compartment and overlap a chromatin environment vulnerable to age-related destabilization. Heterochromatin and H3K9me3 are progressively lost during aging [33], and derepression of LINE/L1 and other transposable elements accompanies neuronal decline in the aging brain [34] and can promote pro-inflammatory signaling in senescent cells [35]. In the neurodegenerative brain, loss of heterochromatin-mediated repeat silencing has been implicated in disease, including tau-induced transposable element dysregulation and neuronal death in tauopathy [36]. Thus, the enrichment of high local SNP-explained VMRs in repeat-associated heterochromatin raises the possibility that inherited variation contributes to individual differences in methylation states within genomic compartments important for long-term epigenomic stability. This positions the genetically anchored methylome not as inert distal sequence, but as a candidate substrate through which population-relevant genetic variation may influence repeat-rich chromatin states relevant to brain aging and neurodegenerative vulnerability.

A small subset of high-SNP intergenic VMRs overlapped ABC-defined enhancer–promoter links and converged on immune-related target genes, including MHC class II loci. This signal was notable because most high-SNP intergenic VMRs did not overlap ABC-defined enhancers, yet the mappable subset was enriched for antigen presentation, T cell activation, and autoimmune disease pathways. In brain tissue, MHC class II expression is concentrated primarily in microglia [37], and genetic variation in the HLA region has been implicated in neurodegenerative, neuropsychiatric, and neuroimmune phenotypes [38,39,40,41,42]. The HLA region also contains strong cis-meQTLs with complex haplotypic structure [43]. These observations suggest that, although the dominant high-SNP VMR architecture is distal and closed, a minority of genetically anchored intergenic methylation sites may connect to immune regulatory programs through enhancer-linked targets.

Low-SNP VMRs showed a contrasting regulatory profile. They were more gene-proximal, more enriched near promoters, enhancers, and transcribed features, and showed reduced cross-region concordance in estimated 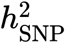. These patterns suggest that low-SNP VMRs capture methylation variation more closely coupled to local tissue regulatory state and less consistently explained by shared local SNP effects. However, environmental and clinical metadata analyses did not support a simple one-to-one mapping between the low-SNP class and environmental exposure. Instead, metadata-associated VMRs were structured by exposure type, brain region, and donor group. Toxicology-defined substance use variables, smoking-related measures, educational attainment, marital status, and trauma history showed context-specific enrichment patterns, with some signals attenuating or reversing in combined-cohort analyses. Multivariate dRFE analyses provided the clearest reproducible signal for educational attainment, although classification performance was modest. At the transcriptional-feature level, environmental convergence was more selective: rank-based and VMR-level tests identified recurrent substance use and psychosocial signals, whereras thresholded overlap enrichment was limited. Thus, low-SNP VMRs appear to mark a more regulatory and context-sensitive methylation compartment, whereas environmental responsiveness is distributed across specific exposures, donor groups, and brain contexts rather than captured by a single VMR class.

Several limitations should guide interpretation. First, discovery sample sizes were modest, although simulations supported elastic-net classification in this range. Sample size also differed across brain regions, with the caudate larger than the DLPFC and hippocampus in both BA and WA donors. Differences in regional power may therefore contribute to the stronger and more reproducible caudate signals observed across the regulatory-gradient, chromatin, and local regulatory-association analyses. Larger region-matched cohorts will be needed to determine the extent to which these regional differences reflect power, measurement stability, or biological architecture. However, technical, cellular, and biological explanations are not mutually exclusive. The caudate has a prominent medium spiny neuron component, whereas the DLPFC and hippocampus contain more heterogeneous mixtures of excitatory neurons, inhibitory neurons, and glia. This cellular structure could contribute to more stable bulk methylation estimates and greater power to detect local SNP contributions. At the same time, human brain methylation and chromatin maps show pronounced region-specific organization, including distinct basal ganglia and striatal epigenomic architectures [25,26,44]. This context is biologically relevant because striatal chromatin regulation is coupled with dopamine-responsive signaling and medium spiny neuron identity [45,46] and caudate molecular and dopaminergic phenotypes are implicated in schizophrenia biology [47,48]. Accordingly, stronger repeat and repressive chromatin signals in the caudate may reflect a combination of greater detectability and region-specific regulatory architecture. Because methylation phenotypes were residualized for age, sex, and schizophrenia diagnosis and trait-level S-LDSC signals were exploratory and did not survive multiple-testing correction, these results are consistent with disease-relevant regional biology but should not be interpreted as evidence for schizophrenia-specific methylation associations. Cell-type-resolved methylation profiling will be needed to disentangle these effects.

Second, ancestry-related differences in LD structure may affect the detectability of local SNP contributions to methylation [49,50,51,52]. Prediction-based 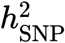 estimation depends on how well observed SNPs tag methylation-influencing variants; therefore, shorter LD and greater haplotypic diversity can reduce prediction confidence at fixed sample sizes and marker density. As a result, some genetically influenced VMRs may be assigned to low-SNP or low-prediction categories. The reported 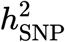 estimates should therefore be interpreted as prediction-based rankings of detectable local SNP contribution under the available sample size, marker density, and LD structure, rather than definitive biological labels for each locus. Third, the available environmental, toxicological, sociodemographic, and clinical variables are broad metadata proxies and cannot resolve exposure timing, duration, or mechanism. Fourth, postmortem brain tissue captures a cumulative molecular endpoint and cannot establish whether methylation differences are causes, consequences, or correlates of lifetime exposures or disease-related processes. Finally, VMR-level analysis improves power but does not resolve CpG-level heterogeneity within regions.

Despite these limitations, our findings establish 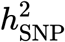 as an organizing axis of the human brain methylome in an underrepresented BA cohort. They distinguish a distal, repressive, repeat-rich methylation architecture with detectable local genetic anchoring—concentrated in compartments governing heterochromatin maintenance and repeat silencing with relevance to neurodegenerative risk—from gene-proximal methylation variation that is less explained by nearby SNPs and more selectively patterned by environmental and clinical metadata. Resolving this architecture in an ancestrally diverse population advances the interpretation of brain epigenomic mechanisms in groups that have been historically underrepresented in genomic studies yet experience inequities in neuropsychiatric and neurodegenerative disease.

## Methods

### Ethical considerations

All research described in this study complies with relevant ethical regulations. All specimens were obtained with oral informed consent in the original studies [13,14]. Specifically, next-of-kin informed consent was obtained under Protocol no. 12-24 from the Department of Health and Mental Hygiene for the Offce of the Chief Medical Examiner for the State of Maryland, and Protocol no. 20111080 from the Western Institutional Review Board for the Offces of the Chief Medical Examiner for Kalamazoo, Michigan; the University of North Dakota in Grand Forks, North Dakota; and Santa Clara County, California.

Original studies obtained samples from the Clinical Brain Disorder Branch at the National Institute of Mental Health (NIMH) from the Northern Virginia and District of Columbia Medical Examiners’ Offces, following NIH institutional review board guidelines (protocol no. 90-M-0142). The Institutional Review Board of the University of Maryland and the State of Maryland approved the study protocols for the collection of these brain tissues [13,14]. Further details on case selection, curation, diagnosis, anatomical localization, and dissection are available in previous publications from the BrainSEQ Consortium [13,14].

### Sample selection and cohort characteristics

We analyzed postmortem brain samples from the BrainSEQ Consortium [13,14,15]. For the discovery cohort, we selected adult case-control samples from the DLPFC (n = 111) [13], hippocampus (n = 116) [13], and caudate nucleus (n = 153) [14]. We included samples that met three criteria: BrainSEQ metadata identified the donor as BA, TOPMed-imputed genotypes were available, and the donor was older than 17 years. The discovery cohort included 380 samples from 168 unique subjects (39% female, 59% controls, mean age 49.6 ± 13.9 years).

For replication analyses, we expanded the dataset by adding study-identified non-Hispanic WA donors from the same BrainSEQ resource. These donors contributed 251 additional samples from 142 unique subjects across the DLPFC (n = 55), hippocampus (n = 60), and caudate nucleus (n = 129). The combined multi-ancestry cohort therefore included 624 samples from 307 unique subjects (34% female, 57% controls, mean age 50 ± 15.2 years). **Tables 1** and **S4** summarize sample characteristics by brain region. **Fig. S5** summarizes the genetic ancestry and population structure of the multi-ancestry cohort.

For these analyses, we treated study-identified race as a proxy for differential exposure to socially patterned environmental conditions, including structural inequities such as residential segregation, unequal access to education and healthcare, and chronic stress, rather than as a biological or genetic category [53].

### BrainSEQ Consortium genotype imputation

We obtained analysis-ready genotype data from the BrainSEQ Consortium [54] through the database of Genotypes and Phenotypes (dbGaP; accession no. phs000979.v3.p2). BrainSEQ previously generated and imputed these data using established procedures [55,56]. Briefly, the consortium genotyped samples on four Illumina microarray platforms (HumanHap650, Human1M, HumanOmni2.5, and HumanOmni5-Quad BeadChips), harmonized data within microarray type, and removed rare variants (minor allele frequency [MAF] < 0.005) and variants failing Hardy–Weinberg equilibrium (P < 1 × 10^−6^). The consortium then converted genomic coordinates from hg19 to hg38 using LiftOver [57] and imputed genotypes on the TOPMed Imputation Server using TOPMed Freeze 8 (GRCh38) reference panels (dbGaP accession number phs001293) [58].

For this study, we applied additional post-imputation quality control and retained variants with imputation *R*^2^ ≥ 0.8, MAF ≥ 0.05, missingness ≤ 0.1, and Hardy–Weinberg equilibrium P 1 × 10^−10^. After filtering, we retained 6,225,756 common variants in BA donors and 6,097,532 common variants in WA donors.

### BrainSEQ Consortium DNA methylation data processing

We downloaded processed CpG-level DNAm data from our previous BrainSEQ study [54] from its GitHub repository (https://github.com/LieberInstitute/aanri_phase1). That study generated WGBS methylation data using a previously described preprocessing pipeline. Briefly, the original pipeline performed raw read quality control with FastQC, removed adapters with TrimGalore [59], aligned reads to hg38 (GRCh38.p12) with Arioc [60], removed duplicate reads with SAMBLASTER [61], filtered reads by mapping quality using SAMtools (v.1.9) [62], and estimated DNAm levels with the Bismark methylation extractor [63]. Additional details of the original WGBS processing workflow are described in our previous study [54].

### DNA methylation quality control

Quality control of the downloaded WGBS methylation data was performed in R (v4.4.3) using the `bsseq` package (v.1.42.0), applied separately to the BA-only discovery cohort and the combined BA + WA replication cohort across three brain regions (caudate nucleus, DLPFC, and hippocampus). We first smoothed methylation levels using `BSmooth`. Low-coverage CpG sites (fewer than five reads in 80% of samples within each brain region), CpG sites overlapping known C/T SNPs, and sites overlapping ENCODE blacklist regions from the `hg38.Kundaje.GRCh38_unified_Excludable` resource available through `AnnotationHub` (v.3.14.0) were excluded.

Sample-level quality was assessed by computing Spearman correlations among all pairs of sample methylation profiles; samples with mean pairwise correlation more than two SDs below the cohort mean were identified as outliers and removed from downstream analysis. To characterize residual technical and biological sources of variation, we performed principal component analysis (PCA) on CpG sites with a minimum per-sample coverage of 10 reads and non-zero variance across at least 80% of samples. The top 20 principal components (PCs) were extracted and their association with sample-level covariates—age at death, post-mortem interval (PMI), the top three SNP-based PCs (snpPC1– snpPC3), brain pH, and age of schizophrenia onset—was assessed using Spearman’s rank correlation. Associations with FDR-adjusted *P* < 0.05 were considered significant; those with *P* < 0.10 were retained as trends. Covariate associations are shown in **Fig. S4**.

### Variably methylated region analysis

For genome-wide heritability analyses, we defined VMRs as genomic regions exhibiting high inter-individual variability in DNAm levels. After quality control (see *DNA methylation quality control*), we quantified DNAm at CpG sites separately in the DLPFC (21,115,720 CpGs), hippocampus (21,384,652 CpGs), and caudate nucleus (24,437,763 CpGs). For each brain region, we first calculated the SD of methylation levels across individuals and retained the top one million most variable genome-wide CpG sites for residualization. Methylation levels at these sites were then regressed on the top three SNP PCs to account for genetic similarity. Using the resulting residual matrix, we performed PCA and regressed out the top five methylation PCs at all CpG sites to reduce batch and other technical effects. We next calculated the variance of the residualized methylation levels and retained the top 1% most variable CpG sites on autosomes and chromosome X (n = 244,367 caudate; 213,845 hippocampus; 211,168 DLPFC) for VMR definition. The number of top 1% variable CpGs per chromosome is detailed in **Table S20**.

We identified VMRs using the `regionFinder3` function from the `bsseq` package (v.1.42.0), with `maxGap = 1000` and a minimum of five CpGs per region. VMR genomic coordinates were recorded in BED format for each chromosome. We then calculated region-level DNAm values for each VMR using `getMeth` with `what = “perRegion”`. To prepare downstream heritability analyses, we stored VMR-level methylation phenotypes in a tab-delimited `.phen` file using the `write_phen` function from the `genio` package (v.1.1.2). Finally, SNPs were extracted within ±500 kb of each VMR using PLINK 2.0 [64] and stored genotype data in PLINK binary format.

We separately extracted VMRs in the multi-ancestry replication cohort. Using the same pipeline for the discovery cohort, we identified 31,662 VMRs for the combined dataset, including 11,457 in the caudate, 9,590 in the DLPFC, and 9,287 in the hippocampus. However, to account for LD differences between populations, we extracted SNPs separately for each population within a larger window (±1 Mb) around each VMR.

In both cohorts, we compared VMR interval length and number of SNPs across brain regions. We fit separate linear mixed-effects models using `lme4` (v.1.1-37) with interval length (*log*_10_ transformed) or number of SNPs as the target variable. We adjusted for the number of samples and included chromosome as a random effect. Pairwise contrasts were evaluated using estimated marginal means with Bonferroni correction, implemented in `emmeans` (v.2.0.2).

### Simulated genotype and phenotype data generation

We generated simulated genotype and phenotype data (**Fig. S1A**) to benchmark the performance of GREML-LDMS and boosting elastic-net regression across seven sample sizes: 100, 150, 200, 250, 500, 1,000, and 5,000 individuals. For each simulation, we generated 1,000 phenotypes and assigned each phenotype to a single autosomal locus with a cis window of ±500 kb (**Fig. S1A**). We designated 25% of phenotypes as high-SNP, assigned one to five causal SNPs to each high-SNP phenotype, and drew narrow-sense heritability values from a uniform distribution between 0.1 and 0.8. For the remaining phenotypes, we drew heritability values between 0 and 0.099 and treated these as low-SNP for downstream classification.

For each locus, we simulated genotypes under an LD structure defined by a Toeplitz correlation matrix with exponential decay (*ρ* = 0.8). We sampled between 150 and 5,000 SNPs per region and generated genotype matrices that approximately satisfied Hardy–Weinberg equilibrium by thresholding multivariate normal draws according to allele-frequency-based cutoffs.

We computed phenotype values as the sum of genetic and environmental components. For high-SNP phenotypes, we calculated the genetic component as a weighted sum of the causal SNP genotypes, with SNP effect sizes drawn from a normal distribution. We then rescaled the genetic component to match the target heritability and added environmental noise so that total phenotypic variance equaled one. We recorded SNP-to-phenotype mappings, simulated heritability values, and genomic coordinates in a tab-delimited summary table for downstream analysis.

### Estimation of SNP-based heritability

#### Genome-wide Complex Trait Analysis

We estimated 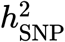 in the simulated genotype–phenotype data using GREML-LDMS as implemented in Genome-wide Complex Trait Analysis (v. 1.94.4) [65]. We applied this analysis at each simulated sample size (n = 100, 150, 200, 250, 500, 1,000, and 5,000). We calculated segment-based LD scores using the default 200 kb window, stratified SNPs into four quartiles according to LD score, and constructed one genetic relationship matrix per quartile. We then fit a multi-component REML model to estimate total 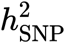 and the variance explained by each LD-stratified SNP group.

#### Boosting elastic-net regression

We estimated local 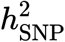 using a custom boosting-based elastic-net framework, building on prior work in boosting-based heritability estimation and elastic-net penalized regression for high-dimensional, correlated genetic predictors [16,17,18,19]. We applied this framework to both simulated phenotypes and VMR-level DNAm phenotypes. For each phenotype, we analyzed SNPs from the corresponding cis window. For real data, we first residualized methylation phenotypes for age, sex, and schizophrenia case-control status using linear regression. Centering and scaling were then performed on these adjusted residuals. Because schizophrenia cases comprised approximately half of the donor cohort, this upfront residualization step was critical to prevent diagnostic status from confounding subsequent VMR-level models.

We loaded genotype data from PLINK binary files using `bigsnpr` (v.1.12.18) [66], removed SNPs with near-zero variance (variance ≤ 10^−6^), and imputed missing genotypes by mode imputation. To reduce redundancy due to LD, we ranked SNPs by their absolute marginal association with the phenotype and performed clumping with `bigsnpr::snp_clumping` using the package defaults (*r*^2^ = 0.2, 500 kb window).

After clumping, we fit an iterative boosting procedure. At each iteration, we selected up to 1000 SNPs with the strongest absolute marginal association with the current residuals and fit an elastic net model using `big_spLinReg` with five-fold cross-validation over *α* ∈ {0.05, 0.10, …, 1.00}. We then predicted the batch contribution, updated the residuals, and accumulated SNP effect sizes across iterations. We ran at most 100 iterations and applied early stopping when the SD of the last five incremental heritability estimates fell below 10^−4^.

We defined the incremental contribution at iteration *t* as Eq. 1:

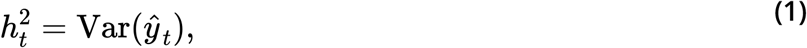

where *ŷ*_*t*_ is the phenotype contribution predicted from the SNP batch selected at iteration *t*. After boosting, we calculated the final unscaled local 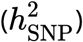 as Eq. 2:

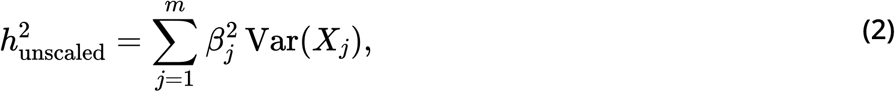

where *β*_*j*_ is the accumulated effect size for SNP *j*, and Var (*X*_*j*_) is the empirical variance of genotype *j* after clumping.

To summarize predictive fit, we refit the clumped SNP set using ridge regression (`cv.glmnet` [18,67,68], *α* = 0) with five-fold cross-validation to select the penalty parameter and calculated predictive fit as Eq. 3 at `lambda.min` :

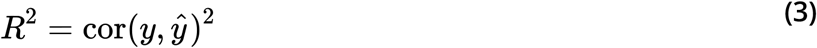

Since SNP ranking and clumping preceded cross-validation, this *R*^2^ represents an internal classification metric, not a true out-of-sample prediction estimate.

### Classification of VMRs into SNP-explained classes

Using the summary statistics from GREML-LDMS and elastic net applied to simulated and real data, we classified phenotypes into three SNP-explained classes based on estimated 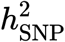 and model prediction confidence:

1. High local SNP-explained sites (high-SNP): 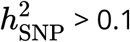, FDR < 0.05 (GREML-LDMS) or *r*^2^ > 0.3 (elastic net))
2. Low local SNP-explained sites (low-SNP): 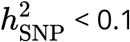, FDR < 0.05 (GREML-LDMS) or *r*^2^ > 0.3 (elastic net)
3. Low-prediction sites: FDR > 0.05 (GREML-LDMS) or *r*^2^ ≤ 0.3 (elastic net).

In the multi-ancestry replication cohort, we estimated 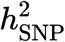 separately for each population using elastic-net regression as described in `Estimation of SNP-based heritability`. This resulted in 13,457 high-SNP sites, 16,534 low-SNP sites, and 268 low-prediction sites for BA and 22,159 high-SNP sites, 7,771 low-SNP sites, and 271 low-prediction sites for WA. Of these, 16,476 VMRs were categorized into the same SNP-explained class across populations (7,304 caudate, 4,145 DLPFC, and 5,027 hippocampus). This set of matched VMRs was used for downstream functional, clinical, and environmental analyses.

To quantify cross-donor-group reproducibility of local genetic architecture, we computed Spearman’s *ρ* between BA and WA 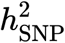 estimates within each SNP-explained class and brain region, restricted to the matched VMRs above. Two-sided *P*-values were obtained from the asymptotic Spearman test (`cor.test`, R v.4.4.3) and concordance scatter plots were generated with `ggscatter` from `ggpubr` (v.0.6.1; **Fig. S10**).

### Sensitivity analysis of VMR classification threshold

To assess the robustness of VMR classification to the choice of prediction confidence threshold, we performed a sensitivity analysis using a more stringent elastic-net performance cutoff. In the primary analysis, VMRs were classified using *r*^2^ ≥ 0.3 to distinguish confidently predicted loci from low-prediction loci. For sensitivity analyses, we repeated the full classification procedure using a stricter threshold of *r*^2^ ≥ 0.75, while retaining the same 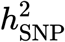 cutoff used to define high-SNP and low-SNP VMRs.

At each threshold, we assigned VMRs to high-SNP, low-SNP, or low-prediction classes and compared class membership across thresholds within each brain region and across all regions combined. We quantified concordance by measuring retention, reclassification, and threshold-specific VMR membership, with particular emphasis on VMRs that shifted from confidently classified to low-prediction under the more stringent criterion. We visualized these patterns using overlap plots and Sankey diagrams.

We applied the same framework to the replication cohort, analyzing BA and WA donors separately. We then compared threshold-dependent classification patterns within each population and examined how VMRs classified as low-SNP in one population were classified in the other (Figs. S7A, S7B, and S7C). Statistical significance was assessed using Fisher’s exact tests with FDR correction.

### Brain regions overlap analysis

We quantified the overlap of VMRs within each SNP-explained class across the caudate nucleus, DLPFC, and hippocampus using `bedtools intersect` (v.2.31.1) [69]. We set the `-F=0.25` option flag to specify the minimum fraction of overlap between VMRs. Specifically, tissue-specific VMRs were defined as regions in which the overlap with the other two brain regions did not exceed 25% of the VMR length. Pairwise shared VMRs were defined as VMRs from one tissue that overlapped VMRs of another tissue by at least 25% of the VMR length from the first tissue. To identify VMRs shared across all brain regions, we first filtered for two brain regions using the `-F` option flag and subsequently examined the overlap of this pre-filtered VMR set with the final brain region using the same overlap threshold. For pairwise intersections, the `-wo` option flag was used to extract the length of overlapping base pairs. Percent overlap was calculated independently for each VMR using *overlap*_*length*_ */ VMR*_*length*_, resulting in two estimates per pair. Reciprocal overlap was recorded as the minimum of the two values. In addition to percent overlap, we quantified similarity using the Jaccard index for each pairwise overlap using `bedtools jaccard`. Summaries of discordance rates and directional class shifts across tissue pairs are reported in Tables S6 and S7.

We visualized sharing of VMRs across brain regions using UpSet plots generated through UpSetR (v.1.4.0). UpSet plots were generated for high-SNP sites, low-SNP sites, low-prediction sites, and all sites. We also analyzed the correlation of overlap between sites (high-SNP, low-SNP, low-prediction, all) across brain regions using Spearman’s correlation tests. Pairwise significance testing was performed using Fisher’s exact tests with FDR correction. We leveraged Monte Carlo simulations to test significance for VMRs shared across all three brain regions. Finally, we generated pairwise correlation plots for visualization using `ggplot2` (v.4.0.1).

### Functional enrichment and genomic annotation analysis

We performed functional enrichment analysis on high-SNP, low-SNP, and low-prediction VMRs to characterize the biological context of genetically and non-genetically patterned methylation regions. For each brain region and SNP-explained class, genomic regions were analyzed with `rGREAT` (v.2.8.0) [70] using Gene Ontology Biological Process (GO:BP), Gene Ontology Molecular Function (GO:MF) [71,72], KEGG [73], UniProt [74], Reactome [75], and the MSigDB C7 ImmuneSigDB collection [76]. For each analysis, we used all VMRs identified in the corresponding brain region as the background universe. We extracted significantly enriched pathways using `getEnrichmentTable` and considered terms with FDR < 0.05 significant.

We also evaluated genomic feature enrichment for each VMR class. We generated GRCh38 annotations using `annotatr` (v. 1.32.0), including basic gene features, intergenic regions, CpG islands and related features, and FANTOM enhancers (`hg38_basicgenes`, `hg38_genes_intergenic`, `hg38_cpgs`, `hg38_enhancers_fantom`). We then tested whether overlap with each annotation class differed across high-SNP, low-SNP, and low-prediction VMRs within each brain region using Fisher’s exact tests, with FDR correction applied across annotation categories.

### Continuous heritability–annotation association analysis

To test whether genomic annotation membership varied with continuous 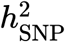 estimations beyond the three-class framework, we restricted analysis to high-confidence VMRs (*r*^2^ > 0.3) and fit logistic regression models separately for each annotation and brain region according to Eq. 4:

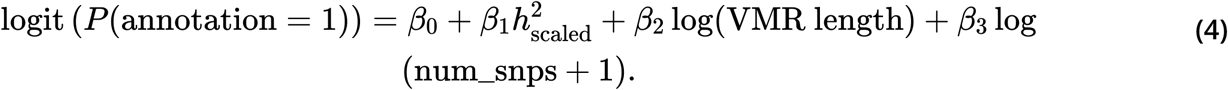

We defined 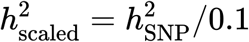 so that ORs correspond to a 0.1-unit increase in 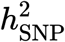. To allow for non-linear relationships, we additionally fit natural cubic spline models with three degrees of freedom and compared these with covariate-only null models using likelihood ratio tests. 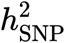 quintiles (Q1– Q5) were constructed per brain region on confidently classified VMRs (*r*^2^ > 0.3) using empirical quintile cutpoints. In the matched multi-ancestry cohort, quintiles were additionally constructed separately for the BA and WA donor groups to expose population-specific architecture effects. Concordant and discordant donor-group-specific quintiles are summarized in (**Fig. S11C**). The same per-region (and per-donor-group) quintile assignments were reused for all downstream quintile-stratified analyses (open chromatin, repressive chromatin, repeat elements, S-LDSC, and local regulatory associations); per-region cutpoints are reported in the corresponding figure legends. We calculated annotation fractions across 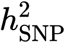 quintiles and estimated 95% Agresti–Coull CIs. We adjusted all *P*-values for multiple testing using FDR correction across annotations and tissues.

### Open chromatin regulatory element enrichment

To evaluate whether high-SNP intergenic VMRs overlap brain cell-type-specific regulatory elements, we intersected intergenic VMRs (`hg38_genes_intergenic`) with single-nucleus ATAC-seq peak sets from the PsychENCODE BrainScope resource [22]. Peak sets were analyzed for seven brain cell types —excitatory neurons, inhibitory neurons, astrocytes, oligodendrocytes, oligodendrocyte precursor cells (OPCs), microglia, endothelial cells—and for a union of peaks across all cell types. We computed overlaps using `GenomicRanges::findOverlaps` (v.1.58.0) [77].

For categorical analyses, we compared high-SNP and low-SNP intergenic VMRs by open chromatin overlap status within each tissue and cell type using Fisher’s exact tests, followed by FDR correction across comparisons. For continuous analyses, we modeled peak overlap as a function of estimated 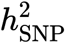 using logistic regression (Eq. 5):

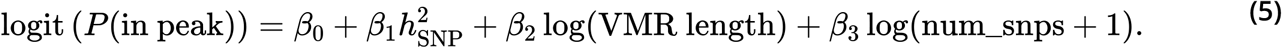

We also compared the top 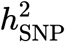 quintile (Q5) with the lower four quintiles (Q1–Q4) using Fisher’s exact tests.

To assign putative target genes, we intersected VMR coordinates with ABC enhancer–promoter links derived from PsychENCODE prefrontal cortex data. For each VMR–gene link, we recorded the target gene, ABC score, and enhancer-to-TSS distance. We then performed GO:BP and KEGG enrichment analyses separately for high-SNP and low-SNP intergenic VMR-linked genes using `clusterProfiler` (v.4.14.0) [78,79] with `org.Hs.eg.db`, using all genes linked to intergenic VMRs as the background universe. Because the ABC annotations were derived from the prefrontal cortex, we applied them to all three brain regions with that limitation noted.

### Repeat element overlap enrichment

To determine whether high-SNP intergenic VMRs are preferentially associated with transposable element-derived sequence, we intersected intergenic VMRs (`hg38_genes_intergenic`) with repeat element annotations from the RepeatMasker track for GRCh38 (hg38) obtained from the UCSC Genome Browser. Repeat elements were analyzed by class (LINE, SINE, LTR, and DNA transposon) and subfamily (L1, Alu, MIR, ERV1, ERVL, ERVL-MaLR, ERVK, and SVA). VMR–repeat element overlaps were computed using `GenomicRanges::findOverlaps` [77], with a VMR considered to overlap a repeat element if any portion of the intervals intersected.

For categorical enrichment testing, we compared high-SNP and low-SNP intergenic VMRs by repeat overlap status within each tissue and repeat class using Fisher’s exact tests, followed by FDR correction across comparisons. We additionally compared the top 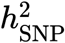 quintile (Q5) against the lower four quintiles (Q1–Q4) and the top decile against the remaining intergenic VMRs using the same approach.

For continuous association analyses, we modeled repeat element overlap as a function of estimated 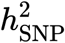 using logistic regression adjusted for VMR length and SNP density, following the same model structure used for open chromatin analysis (Eq. 5).

### Repressive chromatin domain enrichment

To characterize the chromatin state of high-SNP VMRs beyond open chromatin depletion, we overlapped VMR coordinates with repressive histone modification peaks and ChromHMM chromatin state annotations from the NIH Roadmap Epigenomics Consortium [24]. Because Roadmap data were in hg19 coordinates, we lifted VMR intervals from hg38 to hg19 using `rtracklayer::liftOver` (v.1.66.0) with the UCSC hg38-to-hg19 chain file, retaining only VMRs that mapped to a single hg19 interval. We used tissue-matched consolidated epigenomes: E068 (Anterior Caudate), E073 (Dorsolateral Prefrontal Cortex), and E071 (Hippocampus Middle).

For histone modifications, we obtained H3K27me3 and H3K9me3 gappedPeak calls (primary analysis) and broadPeak calls (sensitivity analysis). For chromatin state annotations, we used the ChromHMM 15-state core model and grouped states into five repressive categories: Polycomb (13_ReprPC + 14_ReprPCWk), heterochromatin (9_Het), quiescent (15_Quies), broad repressive union (9_Het + 13_ReprPC + 14_ReprPCWk + 15_Quies), and bivalent (10_TssBiv + 11_BivFlnk + 12_EnhBiv). VMR–peak overlaps were computed using `GenomicRanges::findOverlaps` [77].

We tested enrichment of high-SNP relative to low-SNP VMRs in each repressive annotation using Fisher’s exact tests, applied separately to all VMRs and to the intergenic subset, with FDR correction across all comparisons. To evaluate dose-dependent associations, we modeled repressive mark overlap as a function of continuous 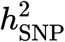 using logistic regression adjusted for VMR length and SNP density (Eq. 6):

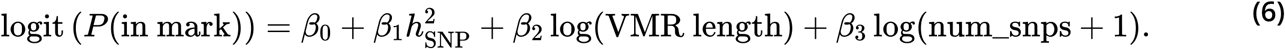

We also compared the top 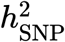 quintile (Q5) and top decile against lower strata within intergenic VMRs using Fisher’s exact tests, with FDR correction. To provide a full chromatin landscape view, we computed the overlap rate between high-SNP and low-SNP VMRs and each of the 15 ChromHMM states.

### Clinical enrichment with stratified LD score regression

We used S-LDSC (v.1.0.1) to test whether GWAS heritability for brain-related traits was enriched in VMR-derived genomic annotations. We tested GWAS summary statistics for smoking initiation, AD, PD, and schizophrenia, restricted to HapMap3 regression SNPs and 1000 Genomes Phase 3 European ancestry reference SNPs (baselineLD v.2.2). To harmonize coordinates between VMRs and the LDSC resources, we lifted VMR intervals from hg38 to hg19 with `pyliftover` (v.0.4.1) using the UCSC `hg38ToHg19.over.chain.gz` chain file, discarding any VMR with unmapped or inverted intervals.

We ran S-LDSC under two complementary annotation schemes. The first was a class-stratified design, in which custom annotation files were generated separately for high-SNP, low-SNP, and low-prediction VMR sets in each brain region across all 22 autosomes. The second was a continuous 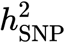 design, in which a single multi-annotation `.annot` file was built encoding membership in each quintile (Q1–Q5) as separate annotations. Quintiles were defined per brain region in the BA discovery cohort and, additionally, per donor group (BA-defined and WA-defined 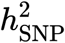 quintiles) in the matched multi-ancestry cohort to expose population-specific genetic-architecture effects. For each scheme, partitioned LD scores were computed against the 1000 Genomes Phase 3 European reference panel and S-LDSC was run for each trait × brain region × annotation set. Coeffcient-scores were converted to two-sided *P*-values and FDR-adjusted with the Benjamini–Hochberg procedure across trait × tissue × quintile cells.

### Environmental association and contribution

Using patient metadata, we initially identified 15 candidate environmental proxies: smoking status, codeine use, morphine use, cocaine use, alcohol use, antipsychotic use, nicotine use, amphetamine use, marital status, sexual abuse history, physical abuse history, other trauma history, military service history, full-scale intelligence quotient (FSIQ), and education level. Because the four trauma items capture related constructs and their missingness partially overlaps, we created a binary composite variable, `any_trauma_hx`, equal to 1 if any of the four items were recorded as positive, 0 if all available items were negative, and missing only when all four were absent. This composite retained more individuals than any single trauma item while reducing the number of correlated tests. The availability of these data is described in Table S13.

To ensure suffcient statistical power and avoid inflating the multiple-testing burden, we applied a data-driven exclusion criterion prior to analysis. For each candidate variable, we computed the proportion of individuals with missing data (after applying all categorical recoding; see below) and excluded variables exceeding 15% individual-level missingness. This threshold excluded FSIQ (>90% missing), antipsychotic use (~19%), and the four trauma/adversity variables (sexual abuse, physical abuse, other trauma, and military service history; each 16–19% missing). The final set of nine variables meeting the missingness criterion comprised: smoking status, codeine use, morphine use, cocaine use, alcohol use, nicotine use, amphetamine use, education level, and marital status.

We correlated VMR methylation levels with these exposures and examined differences within SNP-explained categories (high-SNP, low-SNP, and low-prediction). Education level (less than high school, high school, and more than high school) and marital status (single, married, and previously married) were modeled as three-category outcomes to maximize sample size.

We conducted this analysis using the BA-only discovery cohort and separately for the matched multi-ancestry cohort from BA and WA donors. Separate linear regression models were applied to each VMR as a function of each environmental variable of interest with relevant biological covariates (sex, age, diagnosis, and global ancestry). As sex, age, diagnosis, and global ancestry were corrected for in heritability estimation, they were included here as a negative control. In the combined BA + WA cohort, the DLPFC VMR class counts were substantially skewed (4,264 high-SNP vs. 690 low-SNP; **Table S5**), which can cause complete separation in linear enrichment models and produce extreme or numerically unstable OR estimates. DLPFC combined-cohort enrichment results should therefore be interpreted with caution; where complete separation was detected (infinite maximum-likelihood OR), we report estimates bounded by floating-point precision and flag these in **Table S15**.

We also identified differentially methylated VMRs using matrix regression. We applied separate vectorized linear models relating methylation values as a function of each environmental variable of interest with relevant biological covariates (sex, age, diagnosis, and global ancestry). We subsequently leveraged an Empirical Bayes approach to compute moderated statistics using the `eBayes` function from the `limma` R package (v 3.62.1).

Using both methods, resulting *P*-values for each environmental variable term were adjusted using FDR, and significant VMRs and DMRs at different thresholds (p < 0.05, FDR < 0.1, FDR < 0.05) were saved into separately. Fisher’s exact tests were conducted to identify enrichment patterns across environmental exposures and SNP-explained categories for nominally associated VMRs (p < 0.05) derived from both tests (linear regression and differential methylation).

### Predictive modeling using dynamic recursive feature elimination and dimensionality reduction

To identify VMRs encoding information about environmental exposures and to improve statistical power through dimensionality reduction, we implemented a two-stage prediction pipeline applied to the nine environmental and clinical metadata variables passing the missingness criterion (see *Environmental association and contribution*).

#### Stage 1—dRFE feature selection

We performed supervised classification using dRFE [80]. Analyses were conducted separately for each brain region and VMR SNP-explained category to prevent cross-regional and cross-category information leakage. For each combination, DNAm data were organized into sample-by-feature matrices retaining VMRs present in ≥60% of samples with variance > 1 × 10^−5^. Prediction tasks required ≥40 samples and ≥10 samples per class. Classification used elastic-net-regularized logistic regression (L1 ratio = 0.5, balanced class weights, SAGA optimizer, maximum 5,000 iterations) implemented in dRFEtools (v 0.4.0). Feature selection and evaluation were conducted via stratified five-fold cross-validation. Within each fold, missing values were imputed using the training-set median and features were standardized (zero mean, unit variance) to prevent leakage. dRFE iteratively eliminated 20% of remaining features per step, reserving 20% of the training data for internal validation during elimination. Performance was assessed by ROC-AUC. The optimal feature subset was identified by applying LOWESS smoothing (fraction = 0.3) to retained feature count versus cross-validated AUC, selecting the smoothed maximum [80]. Feature importance was defined as the absolute regression coeffcient (binary outcomes) or the L1-norm across classes (multinomial outcomes), aggregated across folds by averaging. Analyses used dRFEtools, scikit-learn (v 1.8.0) [81], NumPy (v 2.4.2) [82], and pandas (v 2.3.3) [83]. Tasks with a cross-validated ROC-AUC < 0.55 were excluded from downstream analysis.

#### Stage 2—PCA and association testing

For each prediction task retaining an optimal VMR subset from Stage 1, we applied PCA to the selected VMRs to capture the major axes of co-varying methylation while minimizing the number of tests. VMR matrices were imputed (column-wise median) and standardized before PCA. We retained the minimum number of PCs explaining at least 80% of cumulative variance (minimum two PCs).

To test per-PC associations within each brain region, we fitted separate generalized linear models (GLMs) for each PC: binary environmental and clinical metadata outcomes were modeled with logistic regression and ordered categorical outcomes (education and marital status) with cumulative link models using the `ordinal` R package (v.2025.12-29). All models included age, sex, primary diagnosis, and global ancestry proportion as covariates. *P*-values for PC terms were corrected for multiple testing within each task using the Benjamini–Hochberg FDR procedure.

To leverage information across brain regions, we additionally fitted linear mixed models combining data from all three regions. For this cross-tissue analysis, we took the union of dRFE-selected VMRs across regions, applied per-tissue median imputation and standardization, and performed PCA on the combined stacked matrix (retaining ≥80% cumulative variance). The cross-tissue model included brain region as a fixed effect and donor identifier as a random intercept to account for repeated measurements per individual (Eq. 7):

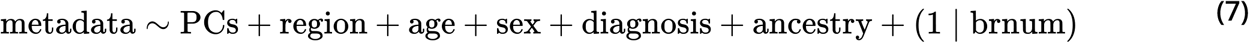

Binary outcomes were fitted with `glmer` (binomial family, bobyqa optimizer) and ordered outcomes with `clmm`, both from the `lme4` and `ordinal` R packages, respectively.

### Bulk RNA-sequencing processing and quantification

We re-mapped BrainSEQ RNA-sequencing reads to the hg38/GRCh38 human reference genome (GENCODE release 47, GRCh38.p14) using the splice-aware STAR aligner (v.2.7.11b) [84] in two-pass mode to improve novel-transcript detection. In the first pass, splice junctions were called across all samples from uniquely and multi-mapped reads, then filtered to remove junctions supported only by multi-mapped reads or by fewer than two uniquely mapped reads; the filtered junction set was incorporated into a second-pass alignment producing coordinate-sorted outputs. Per-sample alignment quality control metrics were collected with RNA-SeQC (v.5.0.4) [85].

We quantified genomic features per sample as follows: gene read counts with featureCounts (v.2.0.8) [86] using paired-end, reverse-stranded parameters; transcript expression (counts and transcripts per million [TPM]) with Salmon (v.1.9.0) [87] for reverse-stranded reads; and alternative-splicing PSI values with SUPPA2 [88] from transcript-level TPM matrices, computing each event type (SE, A3, A5, AF, AL, MX, and RI) per brain region (caudate, DLPFC, and hippocampus). Counts, feature annotation, and quality control metrics were packaged into `RangedSummarizedExperiment` objects [89].

### Expression cell-type proportion estimation

To estimate cell-type composition for use as RNA-level covariates in the local methylation– transcriptional feature association models, we performed reference-based deconvolution of the BrainSEQ bulk RNA-seq data with MuSiC (v.1.0.0) [90], using region-matched Tran et al. single-nucleus RNA-seq references from prior BrainSEQ work [91]. Caudate deconvolution used the Tran et al. nucleus accumbens reference as the nearest available striatal reference, DLPFC used the Tran et al. DLPFC reference, and hippocampus used the Tran et al. hippocampus reference. Reference cell labels were mapped to a harmonized set comprising astrocytes, excitatory neurons, inhibitory neurons, oligodendrocytes, OPCs, microglia, immune cells, and mural cells; caudate additionally retained D1-SPN and D2-SPN medium spiny neurons. Bulk RNA-seq count matrices for the caudate (*n* = 487), DLPFC (*n* = 500), and hippocampus (*n* = 452) were intersected with the single-nucleus reference gene set, and MuSiC was run with `music_prop` to return weighted estimates of per-sample cell-type proportions (**Fig. S12**), which were used as covariates in the local association models described below. Per-region proportions are released as **Supplementary Data 1**.

### Local methylation–transcriptional feature associations and genomic proximity

#### Local association testing

To test whether VMR methylation levels associate with nearby transcriptional features, we linked each VMR to gene expression and alternative splicing targets using a linear regression framework. For gene expression, we applied two complementary linkage strategies: ABC enhancer–promoter links from PsychENCODE prefrontal cortex (the same links used for target gene identification above), yielding VMR–gene pairs constrained to enhancer-overlapping VMRs; and the nearest expressed gene within a 250 kb cis window of the VMR midpoint (nearest gene strategy). For alternative splicing, we paired each VMR with all SUPPA2-defined PSI events within 250 kb of the VMR boundaries. Features with zero variance across samples were excluded prior to association testing within each tissue.

For each VMR–feature pair, we modeled the transcriptional outcome as a linear function of VMR methylation, adjusting for RNA-level covariates (sex, age, primary diagnosis, estimated cell-type proportions, and RNA quality metrics), according to Eq. 8:

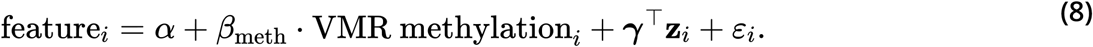

To reduce computational cost for the large nearest gene and PSI pair sets, we precomputed (*X*^⊤^ *X*)^−1^ per VMR and applied batched ordinary least squares estimation across all feature pairs sharing the same covariate structure, reverting to pair-wise fitting when numerical issues arose. We required ≥ 20 samples with complete covariate and outcome data and ≥ 3 distinct VMR methylation values. *P*-values for the methylation coeffcient were adjusted using FDR correction across all tested pairs within each tissue and modality. Significant associations were defined at FDR < 0.05 for expression analyses and FDR < 0.10 for splicing analyses, reflecting the larger multiple-testing burden in the PSI analysis. We summarized results by SNP-explained class as the proportion of VMRs with ≥ 1 significantly linked feature and computed the peak effect size (maximum | *β* |) per VMR.

#### Genomic proximity analysis

To quantify the spatial relationship between VMRs and transcribed features, we computed the distance from each VMR midpoint to the nearest gene TSS and to the nearest tested PSI event boundary using `GenomicRanges::distanceToNearest` [77], using Ensembl GRCh38 protein coding and lncRNA gene annotations and the full set of SUPPA2-tested events per tissue, respectively. We compared high-SNP and low-SNP VMR distance distributions using two-sample Wilcoxon rank-sum tests and Kruskal–Wallis tests across 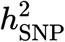 quintiles. Architecture-adjusted logistic regression models – with SNP-explained class as outcome and log-transformed distance, VMR length, and SNP count as predictors – were used to estimate ORs and 95% Wald CIs. FDR correction was applied across tissue and modality comparisons.

#### Environmental convergence

To test whether transcriptional features linked to VMR methylation also associate with environmental exposures, we fit limma vectorized linear models for each transcriptional feature against each of the nine environmental variables meeting the missingness criterion (see *Environmental association and contribution*), adjusting for the same RNA-level covariates used in the local association analysis. Features were classified as environment-linked at FDR 0.10 within each variable. Among features linked exclusively to high-SNP or low-SNP VMRs, we tested whether environment-linked features were overrepresented in one context using Fisher’s exact tests, with FDR correction applied across all environment–tissue combinations.

#### Multi-ancestry replication

We repeated the local association, proximity, and environmental-convergence analyses in the matched multi-ancestry VMR set, stratifying by donor group (BA and WA). Per-donor-group covariate sets were used for the local association and convergence models, while genomic proximity tests operated on the shared matched-VMR coordinates and returned identical distance distributions across donor groups; cohort-specific adjustments in the architecture-adjusted logistic models produced minor BA and WA differences in OR estimates. Summary statistics are reported in **Table S11**, and replication figures appear in **Fig. S19**.

## Acknowledgments

This work was supported by the National Institute on Minority Health and Health Disparities of the National Institutes of Health (NIH) under Award Number R00MD016964 to K.J.M.B. This work is the result of NIH funding, in whole or in part, and is subject to the NIH Public Access Policy. Through acceptance of this federal funding, the NIH has been given a right to make the work publicly available in PubMed Central. This work used the Bridges-2 system at the Pittsburgh Supercomputing Center through allocation BIO250079 from the Advanced Cyberinfrastructure Coordination Ecosystem: Services & Support (ACCESS) program, which is supported by National Science Foundation grants 2138259, 2138286, 2138307, 2137603, and 2138296.

We thank the Lieber Institute for Brain Development (LIBD) and the BrainSEQ Consortium for acquiring, curating, and releasing the data used in this work. We thank the Offces of the Chief Medical Examiner of Washington, D.C., Northern Virginia, Kalamazoo, Michigan, Santa Clara County, California, the State of Maryland, and the University of North Dakota for providing postmortem brain tissue. We also thank the late L.B. Bigelow and the members of the LIBD Molecular Neuropathology Section for their contributions to assembling and curating the clinical and demographic information and maintaining the Human Brain Tissue Repository. We are deeply grateful to the families who donated tissue to advance research on psychiatric and neurological disorders.

## Author contributions

A.B. and K.J.M.B. conceptualized and designed the study. A.B. led formal analysis, visualization, and interpretation of results. E.K.J., N.N.T., and J.H. contributed to statistical analysis and interpretation. K.J.M.B. acquired funding, generated the simulated data, supervised the project, and provided mentorship. A.B. and K.J.M.B. wrote the original draft with input from all authors. All authors reviewed, edited, and approved the final manuscript.

## Web resources

- Genome-wide Complex Trait Analysis (GCTA), https://yanglab.westlake.edu.cn/software/gcta/
- PLINK 2.0, https://www.cog-genomics.org/plink/2.0/
- LiftOver, https://hgdownload.soe.ucsc.edu/goldenPath/hg38/liftOver/
- Stratified LD Score Regression (S-LDSC), https://github.com/bulik/ldsc
- dRFEtools, https://drfetools.readthedocs.io/en/stable/
- NIH Roadmap Epigenomics portal, https://egg2.wustl.edu/roadmap/web_portal/
- UCSC Genome Browser, https://genome.ucsc.edu/
- TOPMed Imputation Server, https://imputation.biodatacatalyst.nhlbi.nih.gov/
- PsychENCODE BrainScope, https://psychencode.synapse.org/

## Data and code availability

Analysis-ready genotype data are available to approved researchers through the database of Genotypes and Phenotypes (dbGaP) under accession no. phs000979.v3.p2. DNA methylation data from BrainSEQ/AANRI Phase 1 are publicly available through GitHub (https://github.com/LieberInstitute/aanri_phase1). Bulk RNA-seq data are available for the DLPFC and hippocampus through the LIBD Globus collections jhpce#bsp2-dlpfc and jhpce#bsp2-hippo and through the BrainSEQ Phase 2 portal (https://eqtl.brainseq.org/phase2/). Bulk RNA-seq data for the caudate nucleus are available through dbGaP accession no. phs003495.v1.p1 and the Caudate eQTL Browser (https://erwinpaquolalab.libd.org/caudate_eqtl/).

For chromatin and regulatory annotation analyses, we used PsychENCODE BrainScope single-nucleus ATAC-seq peaks (http://brainscope.psychencode.org/), repressive histone modification peaks, and ChromHMM chromatin-state annotations from the NIH Roadmap Epigenomics Consortium (https://www.ncbi.nlm.nih.gov/geo/roadmap/epigenomics/?view=matrix) and Activity-by-Contact enhancer–promoter links. For S-LDSC analyses, we used HapMap3 regression SNPs from the LDSC resource (https://data.broadinstitute.org/alkesgroup/LDSCORE/w_hm3.snplist.bz2), 1000 Genomes Project Phase 3 reference data (https://www.internationalgenome.org/data/), and publicly available GWAS summary statistics for smoking initiation (http://genome.psych.umn.edu/research/gscan), Alzheimer’s disease (https://www.ebi.ac.uk/gwas/publications/35379992), Parkinson’s disease (https://www.ebi.ac.uk/gwas/studies/GCST009325), and schizophrenia (https://figshare.com/articles/dataset/scz2022/19426775). Genomic-coordinate conversion, when required, used the hg38ToHg19 LiftOver chain file from the UCSC Genome Browser (https://hgdownload.soe.ucsc.edu/goldenPath/hg38/liftOver/). For cell deconvolution, we used Tran et al. human brain region single-nucleus RNA-seq reference datasets from prior BrainSEQ work, available on GitHub (github.com/LieberInstitute/10xPilot_snRNAseq-human).

Supplementary data generated in this study are available on Zenodo (https://doi.org/10.5281/zenodo.20547606). Analysis code is available at https://github.com/heart-gen/dna-methylation-heritability.

